# BMP-dependent oligoclonal cancer attractor state precedes neural crest fate in melanoma initiation

**DOI:** 10.1101/2024.10.22.618007

**Authors:** Alicia M. McConnell, Maggie H. Chassé, Haley R. Noonan, Jeffrey K. Mito, Julia Barbano, Erika Weiskopf, Irene F. Gosselink, Song Yang, Kyle D. Drake, Chloé S. Baron, Elise Layne, Meera Prasad, Phammela Abarzua, Maxime Grimont, Yung-Ching Kao, Christine G. Lian, Anaïs Eberhardt, Marie Donzel, Katie J. Lee, Iwei Yeh, George F. Murphy, Mitchell S. Stark, Stéphane Dalle, Julie Caramel, Cole Trapnell, Leonard I. Zon

## Abstract

Field cancerization posits that groups of cells harboring oncogenic mutations create a permissive landscape predisposed to malignant transformation. We previously identified rare single cells in BRAF^V600E^;p53^-/-^ zebrafish that reactivate an embryonic neural crest program prior to melanoma initiation. Here, we identify a specific field of BRAF^V600E^;p53^-/-^melanocytes with altered differentiation, morphology and cell-cycle regulatory programs that predates the neural crest activation. Based on single cell analysis, these cells form a cancer precursor zone (CPZ) from which a single clone ultimately stabilizes a neural crest–like state and expands to form melanoma. Using in vivo cellular barcoding combined with single-cell RNA-seq and ATAC-seq, we identify a transcriptionally distinct attractor state specific to oligoclonal CPZs that is modulated by BMP signaling.

Mechanistically, BMP-dependent induction of the transcriptional repressor ID1 sequesters TCF12, thereby inhibiting lineage-specific targets required for maintenance of melanocyte identity and for clonal selection. Single cells from the CPZ initiate neural crest reprogramming and become tumorigenic. Analysis of a large human patient cohort reveals high ID1 expression in precursor melanocytes as early as dysplastic nevi and atypical melanocytic proliferations, implicating ID1 in early human melanomagenesis.

This work identifies BMP signaling and ID1 as early, targetable vulnerabilities with potential for improved diagnosis and prevention of melanoma. Together, these findings uncover a previously unrecognized field effect during melanoma initiation, in which tumors emerge from an oligoclonal attractor-state zone of morphologically distinct yet clinically covert precursor cells with a defined altered transcriptional fate.

---

Melanoma incidence has increased dramatically over the past three decades, yet the earliest stages of melanomagenesis remain poorly defined. Multiple studies report substantial variability among pathologists in the diagnosis of early precursor lesions, underscoring a fundamental gap in our understanding of melanoma initiation(*1-4*). Over 80% of melanomas harbor activating MAPK pathway mutations, most commonly *BRAF*^V600E^, frequently accompanied by tumor suppressor loss-of-function mutations(*5, 6*). However, *BRAF*^V600E^ alone is insufficient to drive melanoma, as 82–100% of benign nevi carry this mutation yet never progress(*7, 8*). Further, subclinical fields of *BRAF*^V600E^ are prevalent in human skin, with enrichment near prior melanoma excisions and benign nevi(*9*). These observations suggest that many potential melanoma precursors remain clinically covert and molecularly undefined.

To investigate the earliest determinants of melanoma initiation, we developed a zebrafish model in which p53^-/-^ melanocytes express BRAF^V600E^. Despite the presence of a widespread cancerized melanocyte field, individual fish develop only 1-3 tumors, with minimal additional mutational burden(*10, 11*). This constrained tumor output mirrors human disease and implies the existence of rate-limiting, non-genetic barriers to malignant transformation. Understanding how genetically primed melanocytes clonally expand may therefore be central to melanoma initiation, yet the clonal dynamics of melanomagenesis remain largely unexplored, in contrast to other tumor types(*12-16*).

We previously demonstrated that individual BRAF^V600E^;p53**^⁻/⁻^** melanocytes undergo neural crest reprogramming to initiate melanoma(*11, 17*). Here, we identify oligoclonal cancer precursor zones (CPZs), a previously unrecognized intermediate state composed of morphologically and transcriptionally aberrant melanocytes. Time-course imaging shows that CPZ formation is necessary for tumor initiation, marking a key transition in melanoma progression. CPZ melanocytes upregulate BMP signaling, and cellular barcoding studies indicate oligo clonal expansion and subsequent clonal dominance and progression toward monoclonal tumors. This process depends on ID1-mediated sequestration of TCF12, which represses differentiation programs and stabilizes a precursor attractor state. Together, our data position CPZs as a critical bottleneck in melanoma initiation and provide a mechanistic framework for how permissive melanocytes transform and reinforce neural crest–like programs to drive melanoma.

## Melanoma forms in oligoclonal cancer precursor zones

To characterize the earliest stages of melanoma formation, we generated *p53* deficient, *crestin:EGFP* zebrafish in which melanocytes express BRAF^V600E^, are marked red by *mitfa:mCherry,* and are amelanotic (referred to as *BRAF;p53*)(*11, 18, 19*). We performed time course imaging and found that prior to neural crest reactivation, there were regions of hundreds of morphologically aberrant melanocytes expressing high levels of *mitfa:mCherry*, which we term the cancer precursor zone (CPZ; Fig. 1A, Supp. Fig. 1A-C). We followed a cohort of fish that exhibit varying kinetics of CPZ formation and neural crest reactivation with an average onset of 11- and 15-weeks post-fertilization (wpf), respectively (Fig. 1B, Supp. Fig. 1D). While on average only 70% of CPZs reactivate *crestin:EGFP* to form a tumor, individual neural crest reactivated *crestin*+ cells within the CPZ inevitably go on to form a melanoma and crestin+ cells do not arise from non-CPZ sites (Fig. 1A,B; Supp. Fig 1B,C). We correlated our imaging with histological analysis of mitfa and crestin across initiation stages, confirming that individual *crestin* positive cells give rise to tumors (Fig. 1C). Further, CPZs form in two additional zebrafish melanoma models (*mcr:BRAF^V600E^* and *mcr:NRAS^Q61R^*) without p53 inactivation, suggesting cancer precursor zones are not specific to BRAF^V600E^ and are not dependent on p53 loss (Supp. Fig. 1E,F).

**Figure 1.**
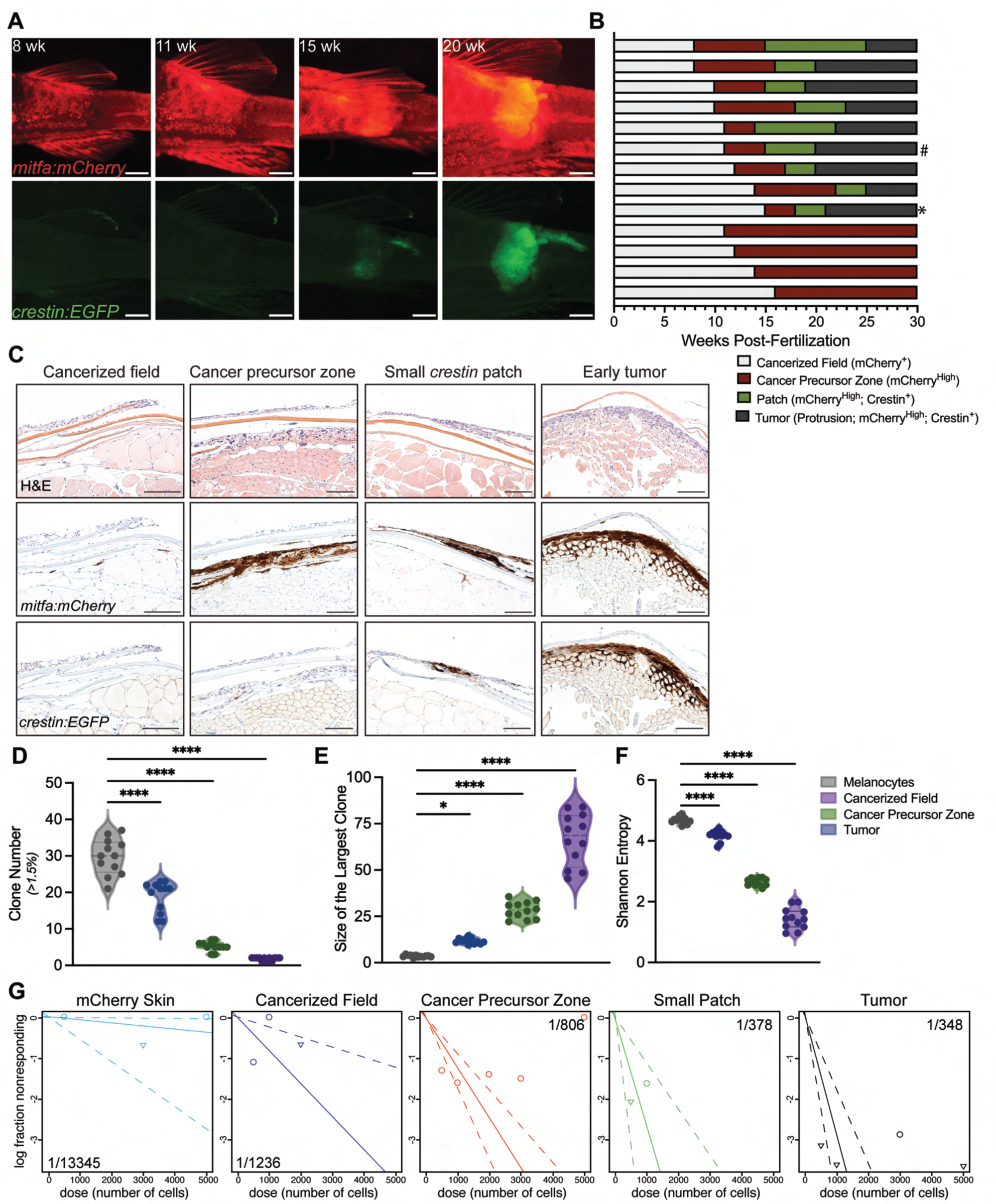
Cancer precursor zone formation precedes neural crest reactivation in zebrafish. A, Melanoma initiation in zebrafish occurs in stages, beginning with the formation of a cancerized field of rescued melanocytes, which expand to form a cancer precursor zone. Neural crest reactivation occurs in cells within the cancer precursor zone and tumor formation follows, with all tumor cells becoming *crestin:EGFP*+. Scale bar: 1 mm. **B,** Swimmers plot of 13 zebrafish during melanoma initiation shows variable onset of CPZ and neural reactivation (# represents fish time course in Fig. 1A; * represents fish time course in Supp. Fig. 1C). **C**, Transverse sections of zebrafish skin stained for mCherry and EGFP at the cancerized field, cancer precursor zone, neural crest reactivation, and tumor formation stages. Scale bars, 100 μm. **D-F,** Violin plot of the total clone number with a minimum clone size of 1.5% (**D**), the largest clone as measured by the clone with the largest percent of total reads (**E**), and the Shannon entropy (**F**) in a cohort of GESTALT barcoded fish (n=12 fish). *, p≤0.05; ****p≤0.0001. **G**, Limit dilution curves at 14 days post-transplant (dpt). Confidence interval estimate is displayed as 1/(engraftment cell frequency).

### Cancer precursor zones are oligoclonal and have aberrant morphology

In line with previous studies, CPZs arise across the entire fish body with more frequent localizations in the head, dorsal fin, and tail base(*20*) with a range of sizes (0.54±0.27mm^2^) and *mitfa:mCherry* intensities (105.6±27.4 a.u.) (Supp. Fig. 2A-C). Despite the presence of oncogenic mutations, the cancerized field melanocytes appear morphologically normal, with proper dendrite formation and spacing throughout the skin (Supp. Fig. 3A, left). However, upon CPZ formation, the melanocytes exhibit aberrant morphology with loss of dendrites (Supp. Fig. 3A, right). Further, histological analysis shows that CPZ melanocytes encompassing *crestin:EGFP*+ cells are highly proliferative with large nuclei (Supp. Fig. 3B,C). Our zebrafish model reveals a novel intermediate step between cancerized field melanocytes and tumor initiation, where morphologically abnormal cancer precursor zone melanocytes encompass the cell undergoing neural crest progenitor state activation as it transforms.

To understand how melanoma clonally evolves, we utilized GESTALT, an in vivo CRISPR-Cas9 based lineage tracing approach, in its first application of melanoma initiation and solid tumor evolution(*21, 22*). We induced two rounds of sgRNA-directed barcoding of the GESTALT cassette at the single cell stage and at 48 hpf, corresponding to the birth of the adult melanocyte stem cell(*23*). We sorted *mitfa:mCherry*+ melanocytes from each melanoma stage and analyzed clone number following DNA sequencing of the GESTALT cassette (Supp. Fig. 4A-D). Healthy melanocytes (29.8±4.9 clones) and cancerized field melanocytes (18.9±4.2 clones) are polyclonal with no evidence of clonal dominance, as indicated by a lack of clones reaching greater than 20% of total reads (three S.D. higher than the average cancerized field clone). CPZ formation is oligoclonal compared to normal skin melanocytes (5.3±1.3 clones) while tumors form from 1-2 clones (1.8±0.5 clones) (Fig. 1D). This marked decrease in clone number was accompanied by a significant increase in the size of the dominant clone (Fig. 1E). Further, the decreased Shannon entropy with a corresponding increase in Gini coefficient and leftward shift in the Lorenz curves as melanoma progresses highlights a decline in clonal diversity in CPZs and the subsequent emergence of a dominant malignant clone (Fig. 1F, Supp. Fig. 4F-J). Together these data indicate melanoma arises through a stepwise clonal bottleneck, with polyclonal melanocytes developing into to oligoclonal CPZs and ultimately a single dominant tumor clone.

To determine if CPZs have malignant potential, we FACS isolated melanocytes from normal skin, the cancerized field, and cancer precursor zones, as well as small *crestin+* patches, or tumors for transplantation (Supp. Fig. 5A). Cells injected under the skin of irradiated recipient fish and a limit dilution assay was performed at 14 days post-transplant. CPZ cells engrafted at about half the rate (0.43x) of *crestin+* cells from small patches or tumors, while normal or cancerized field melanocytes did not engraft (Fig. 1G, Supp. Fig. 5B). Small *crestin* patch cells engrafted at a similar rate to tumor cells, suggesting that neural crest reactivated cells have similar malignant potential. Together, these data demonstrate a substantial clonal selection from the cancerized field melanocytes, providing a set of cells that gain malignant potential and could become tumorigenic. Further clonal restriction occurs from the pool of CPZ clones to form monoclonal tumors.

### Cancer precursor zones are transitory with distinct transcriptional and chromatin states

*BRAF;p53* zebrafish only develop one to three melanomas over their lifetime with minimal additional genetic mutations(*10*), indicating that transcriptional changes are key drivers of melanoma initiation. To explore this, we performed single-cell RNA-seq (scRNA-seq). Control cancerized field, cancer precursor zones, or small *crestin* patches were isolated and run through the 10X Genomics scRNA-seq platform (Fig 2A, Supp. Fig. 6A,B). scRNA-seq module scores for mitfa+/mCherry+ cells were elevated in both CPZ and crestin+ patch populations, whereas crestin+/EGFP+ module scores were high only in crestin+ patch cells, consistent with the dynamics observed in time-course imaging (Fig. 1A, Fig. 2B,C). Dimensional analysis highlights cellular reprogramming from cancerized field melanocytes to crestin+ melanoma with CPZ melanocytes as a transitory population, with an independent analysis using InDrop scRNA-seq showing similar results (Fig. 2D-I, Supp. Fig. 6C-H). CPZ cells express both melanoblast (e.g., *tfap2a*, *tfap2e, runx3)* and neural crest markers (e.g., *foxd3* and *sox10*) highlighting the transitory transcriptome of this population (Fig. 2J). These analyses reveal that the transformation of a terminally differentiated melanocyte occurs through a transcriptional shift through a cancer precursor zone and towards a neural crest progenitor state.

**Figure 2.**
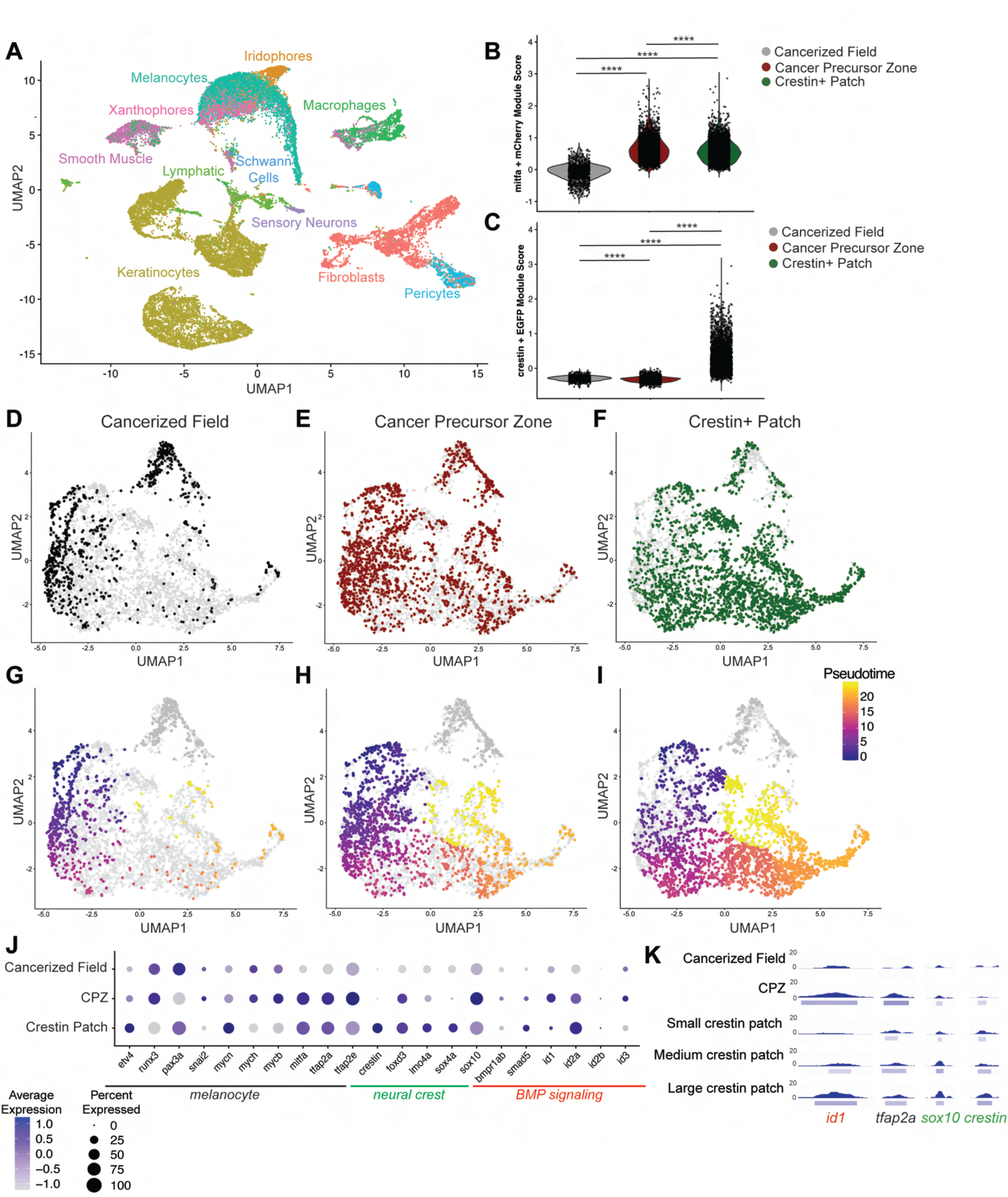
Single cell expression atlas of melanoma initiation. **A**, UMAP of the melanoma initiation atlas resulting from cancerized field (n=2), CPZ (n=4), and patch (n=2)-bearing zebrafish. **B, C.** Violin plot highlighting module enrichment for mitfa and mCherry expression (**B**) or crestin and EGFP expression (**C**) for each cell within the cancerized field, CPZ, or crestin+ stage. **D-F,** UMAP highlighting individual melanoma stages for cancerized field (**D**), CPZ (**E**), and crestin+ (**F**) melanocytes. **G-I,** UMAP of Pseudotime analysis with cancerized field melanocytes set as the root in cancerized field (**G**), CPZ (**H**), and crestin+ (**I**) melanocytes. Dark gray cells are disconnected from the main trajectory. **J**, Differentially expressed genes categorized into melanocyte (black), neural crest (green), and BMP signaling (red). **K**, Open chromatin peaks in promoter or enhancer regions of stage-specific genes.

To verify CPZ cells are a transitory population in melanoma initiation, we performed RNA-seq on FACS isolated *mitfa:mCherry+* cancerized field melanocytes and compared them to *mitfa:mCherry+* CPZ melanocytes and *mitfa:mCherry+/crestin:EGFP*+ cells isolated from small, medium, and large *crestin* patches. CPZ induction did not cause a global impact on gene expression relative to cancerized field with only 502 genes (2.8% [n=17,483], log_2_FC±2, p-value <0.001) altering in expression, indicating this transcriptional switch is specific to progression in melanomagenesis (Supp. Fig. 7A-C).Due to these changes in gene expression, PCA analysis shows that the sorted melanocytes cluster according to stage, with the cancer precursor zone melanocytes overlapping both the cancerized field and *crestin* patch clusters (Supp. Fig. 7D). This suggests that the cancer precursor zone cells represent a transition state between the cancerized field and neural crest reactivated cells.

The distinct gene expression signatures separating melanocytes, cancer precursor zones, and *crestin* positive cells indicates that melanoma initiation occurs through unique transcriptional cell states. Differential expression analysis of both the scRNA-seq and RNA-seq data revealed stage-specific expression signatures (Fig. 2J, Supp. Fig. 8A). CPZ cells have increased expression of BMP signaling (red), as well as neural plate border and pre-migratory neural crest transcription factors (green). These expression profiles were correlated with gene expression at various stages of human melanoma formation from two published datasets (Supp. Fig. 8B,C)(*24, 25*). Together, these findings support a conserved, stage-specific transcriptional trajectory in which melanoma initiation arises from a distinct cellular state that mirrors programs observed during human melanomagenesis.

To dissect the chromatin landscape, we performed ATAC-seq on sorted melanocytes from various stages of initiation. We filtered called peaks for direct overlap or within a 5000 bp distance of annotated genes for differential accessibility analysis (multiple tests corrected with FDR <0.05). Only 999 genes (2.8% [n=35,592 total], log_2_FC±2.0, p-value <0.001) had differentially accessible regions (DAR) in cancer precursor zone melanocytes relative to cancerized field (Supp. Fig. 9A-C). Promoter and enhancer regions associated with differentially expressed genes contained peaks that opened in a stage-specific manner (Fig. 2K, Supp. Fig. 10). Like RNA-seq, ATAC-seq showed reprogramming of specific loci. and further indicate that a transcriptionally driven cancer attractor state is present at the earliest stages of melanoma initiation.

### Cancerized field lacks MAPK activation

Most human melanomas arise de novo, with only a small fraction of nevi progressing to malignancy(*26*). However, 82-100% of benign nevi carry the *BRAF*^V600E^ mutation and activation of the MAPK pathway does not always directly correlate with *BRAF* mutations in many melanomas(*7, 8, 26-29*). Despite the presence of constitutively active BRAF in our zebrafish model, the expression levels of MAPK pathway members and downstream targets are upregulated exclusively in cancer precursor zone and *crestin+* cells (Supp. Fig. 11A). Negative regulators of MAPK, *spry2*, *spry4*, *dusp2*, and *dusp6*, show little to no expression, with corresponding closed chromatin in the cancerized field melanocytes (Supp. Fig. 11A, B). Importantly, ATAC-seq analysis on isolated melanocytes revealed that the enhancer regions around MAPK target genes are not open in cancerized field but become open in cancer precursor zones (Supp Fig. 10B). Furthermore, phospho-ERK immunohistochemistry showed no staining in cancerized field melanocytes, low staining in *mitfa*^med^ CPZs, and high staining in *mitfa*^high^ CPZs (Supp. Fig. 11C). We considered the possibility that the *mitfa*^high^ state increases the amount of BRAF^V600E^ expression due to the *mitfa:BRAF^V600E^*transgene; however, RNA-seq on isolated melanocytes shows no change in BRAF^V600E^ expression between the cancerized field and CPZ stages, only increasing at the patch stage (Supp. Fig. 11D). Together these data indicate that the MAPK pathway is not active until the cancer precursor zone stage and is further activated during tumor initiation.

### ID1 identifies and drives precursor zone formation

Given that cancer precursor zones form prior to, and have a distinct transcriptional state from cancerized field or *crestin* patches, we hypothesized the CPZ state is independently regulated. As CPZs had high expression of BMP pathway members and downstream targets (Fig. 2J), and BMP signaling has an established role in neural crest development(*30-32*), we next interrogated the role of BMP signaling in melanoma initiation. We first confirmed endogenous upregulation of *ID1* in CPZs by RNAscope in situ hybridization(*33*). *ID1* is highly expressed in CPZ melanocytes but not in adjacent cancerized field cells (Supp. Fig. 12). Next, we used the MiniCoopR system to overexpress *ID1*, a constitutively active *SMAD1* (caSMAD), or a dominant negative *BMP* receptor (dnBMPR) in melanocytes(*18*). Activation of BMP signaling (MCR:caSMAD) significantly increased CPZ incidence and subsequent tumor onset, while inhibition of BMP signaling (MCR:dnBMPR) reduced CPZ and tumor incidence (Fig. 3A-D). Taken together, these data support a model in which BMP signaling regulates the emergence of CPZs, establishing a critical early step in melanoma initiation.

**Figure 3.**
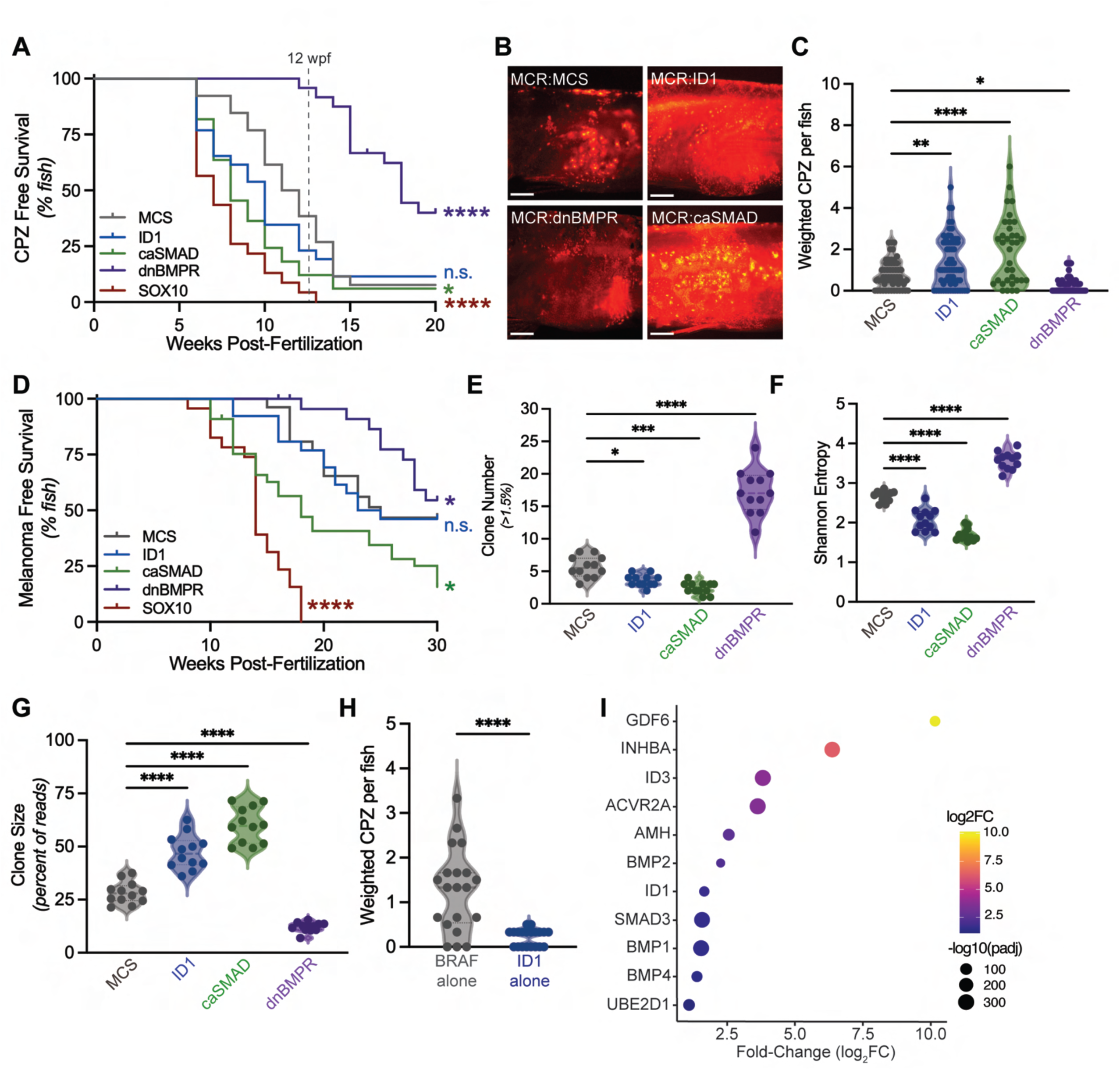
ID1 induces cancer precursor zone formation. A, Kaplan-Meier curve of CPZ incidence following modulation of the BMP pathway compared to MCS (control) and SOX10 (known accelerator). n.s. = not significant; *p≤0.05; ****p≤0.0001; n>15. **B**, Images showing *mitfa:mCherry* fluorescence in zebrafish overexpressing ID1, caSMAD, or dnBMPR compared to an empty MCS control (12 wpf). **B**, Violin plot showing the weighted number of cancer precursor zones per fish. Each dot represents one fish. * *p* ≤ 0.05; ** *p* ≤ 0.01; **** *p* ≤ 0.0001. **D,** Kaplan-Meier curve of tumor incidence following modulation of the BMP pathway compared to MCS (control) and SOX10 (known accelerator). n.s. = not significant; *p≤0.05; ****p≤0.0001; n>15. **E-G,** Violin plot of the total clone number with a minimum clone size of 1.5% (**E**), Shannon entropy (**F**), and the largest clone as measured by the clone with the largest percent of total reads (**G**) in a cohort of GESTALT barcoded fish (n=12 fish). *, p≤0.05; ****p≤0.0001. **H,** Violin plot showing the weighted number of cancer precursor zones per fish in MCR:BRAF^V600E^ relative to MCR:ID1 in wild type fish. Each dot represents one fish. ****p≤0.0001. **I,** Dot plot of enriched BMP marker genes in PMEL(BRAFV600E) human cells relative to PMEL without BRAF(V600E).

We next hypothesized BMP activity modulates clonal expansion in melanomagenesis. Indeed, activation of BMP signaling (MCR:caSMAD) decreased CPZ clone number (MCR:MCS: 5.6±1.6; MCR:ID1: 3.6±0.9; MCR:caSMAD: 2.4±0.9 clones) and Shannon entropy while driving clonal expansion, leading to a shift in the Lorenz curve, without significantly altering the Gini coefficient relative to control as measured by GESTALT cellular barcoding (Fig. 3E-F, Supp. Fig. 13). Conversely, BMP inhibition (MCR:dnBMPR) increased CPZ clone number (17.3±3.4 clones) and Shannon entropy, reduced the dominant clone’s size and lowered the Gini coefficient (Fig. 3E-F, Supp. Fig. 13). Together, these data indicate that BMP activation drives clonal consolidation towards oligoclonality, whereas BMP inhibition promotes a more diverse polyclonal landscape.

While BMP is necessary for CPZ clonal expansion, ID1 overexpression is not sufficient to drive CPZ formation in the absence of BRAF^V600E^ (Fig. 3H). Next, to highlight the role of MAPK signaling in BMP activation, RNA-seq of human primary immortalized melanocytes (PMEL(*34*)) with and without BRAF^V600E^ (PMEL-BRAF) exhibited differential gene expression of known BMP activators and responders highlighting the activation of BMP signaling following BRAF^V600E^ overexpression, as we had observed in the zebrafish (Fig. 3I). Overall, these results provide evidence that the stages of melanoma initiation are independently regulated with BMP signaling driving CPZ initiation and not only clonal selection to tumorigenic cells. Perturbation of signaling leads to a clonal selection and expansion, providing a pool of clones for subsequent tumor formation.

To verify the role of ID1 in human melanoma, we performed immunofluorescence staining on patient samples from four independent cohorts across various melanoma stages from healthy skin to thick melanoma. The percent of melanocytes expressing ID1 and the intensity of ID1 staining was significantly increased in patient samples with atypical melanocyte proliferative (potentially precursor) zones, melanoma in situ, or invasive melanoma compared to those in normal skin (Fig. 4A-D, Supp. Fig. 14). In addition to more cells expressing ID1 as melanoma initiates, cells also have higher ID1 expression (Fig. 4E). The ability of ID1 to initiate CPZs in the zebrafish and the appearance of ID1 in atypical melanocytic hyperplasia fields validate the role for ID1 in melanomagenesis and identify clinically covert ID1+ melanocytic precursor lesions in humans.

**Figure 4.**
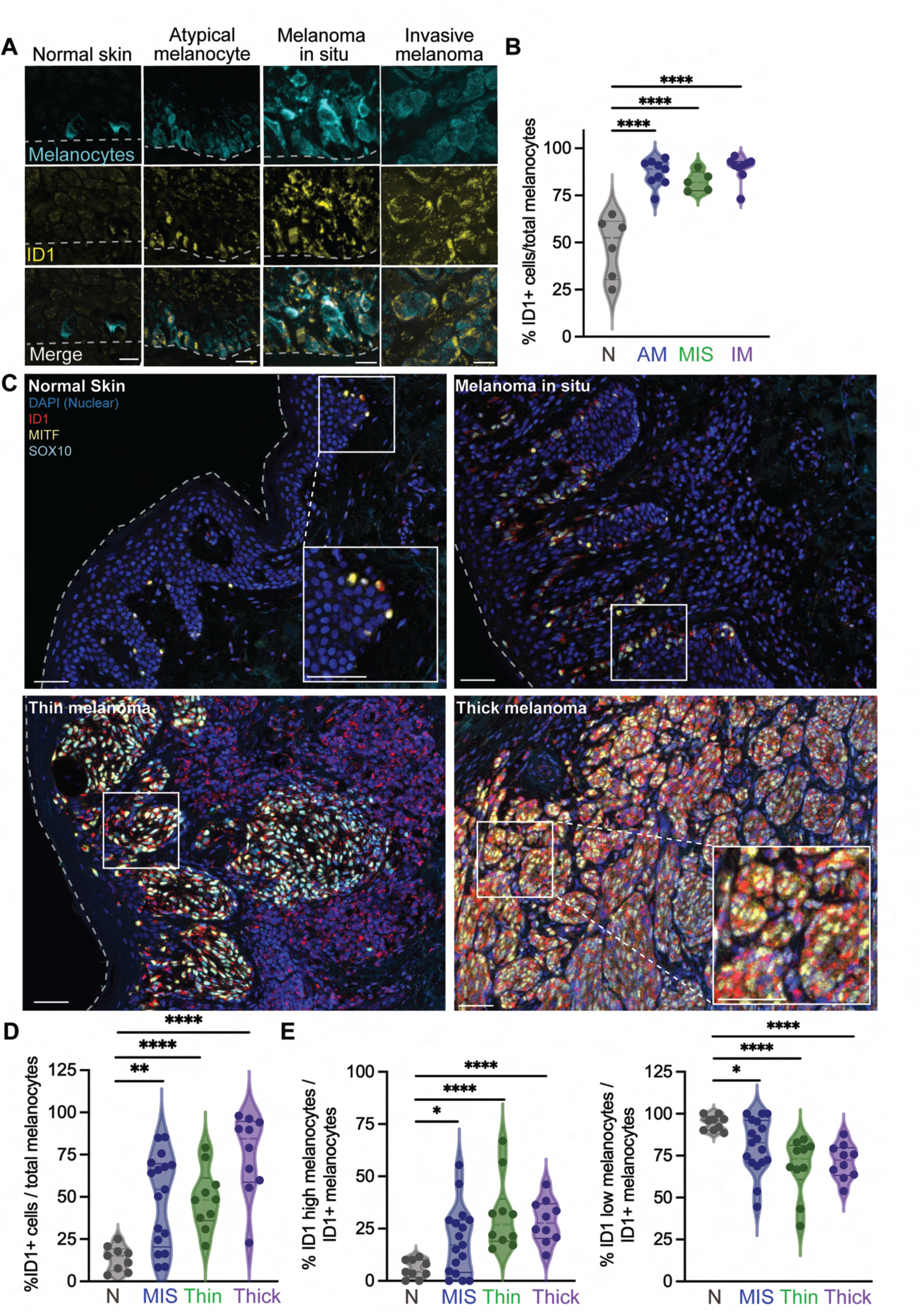
ID1 is upregulated during human melanoma initiation and progression. A, Immunofluorescent staining of melanocytes (MART1) and ID1 in human skin samples. Scale bars, 10 μm. **B**, Percent of melanocytes that are positive for ID1. Each dot represents one patient (n =5-9). **p≤0.01; ****p≤0.0001. **C**, Immunofluorescent staining of melanocytes (SOX10 or MITF, yellow), ID1 (red), and DAPI (blue) in human skin and melanoma samples. Inset highlights regions of double positive melanocytes (MITF/SOX10^+^; ID1^+^). Scale bars, 50 μm. **D,** Percent of melanocytes that are positive for ID1. Each dot represents one patient (n = 9-17). **p≤0.01; ****p≤0.0001. **E,** Percent of melanocytes that are positive for ID1 with high ID1 expression (left) or low ID1 expression (right) across melanoma stages. Each dot represents one patient (n = 9-17). p≤0.05; **p≤0.01; ****p≤0.0001.

### ID1 sequesters TCF12 from chromatin to drive initiation

ID1 is overexpressed in over 20 types of cancer, including melanoma, where strong ID1 expression was significantly associated with increased tumor thickness and decreased patient survival(*35, 36*). ID1 is a transcriptional repressor that prevents basic helix-loop-helix (bHLH) transcription factors from binding to DNA(*37*). We performed IP-MS on A375 melanoma cells overexpressing V5-tagged ID1 or Clover control. The bHLH factor TCF12 was the most significantly pulled down protein with ID1 (Fig. 5A, Supp. Fig. 15A). When performing ATAC-seq on sorted melanocytes isolated from ID1-overexpressing zebrafish, we found significant enrichment of the TCF12 motif (ACAGCTG) under peaks that decreased with ID1 overexpression, indicating ID1 is a repressor of TCF12 signaling (HOMER, p=1e-20). Predicted downstream targets of TCF12 are significantly decreased in CPZs where ID1 is most highly expressed (Supp. Fig. 15B)(*38*). Next, we performed RNA-seq on sorted melanocytes from zebrafish overexpressing ID1 and identified potential downstream targets that were downregulated following ID1-overexpression or in CPZs (Supp. Fig. 15C,D). TCF12 motifs were found under open chromatin peaks in the promoter/enhancer regions associated with these genes, many of which are important for dendritic processes and cell adhesion programs (e.g., *rhogb*, *parvg*, *lamc1*, and *enah*) (Fig. 5B). Together, these results indicate that ID1 promotes melanoma progression by repressing TCF12-mediated transcriptional programs critical for melanocyte differentiation and cell proliferation.

**Figure 5.**
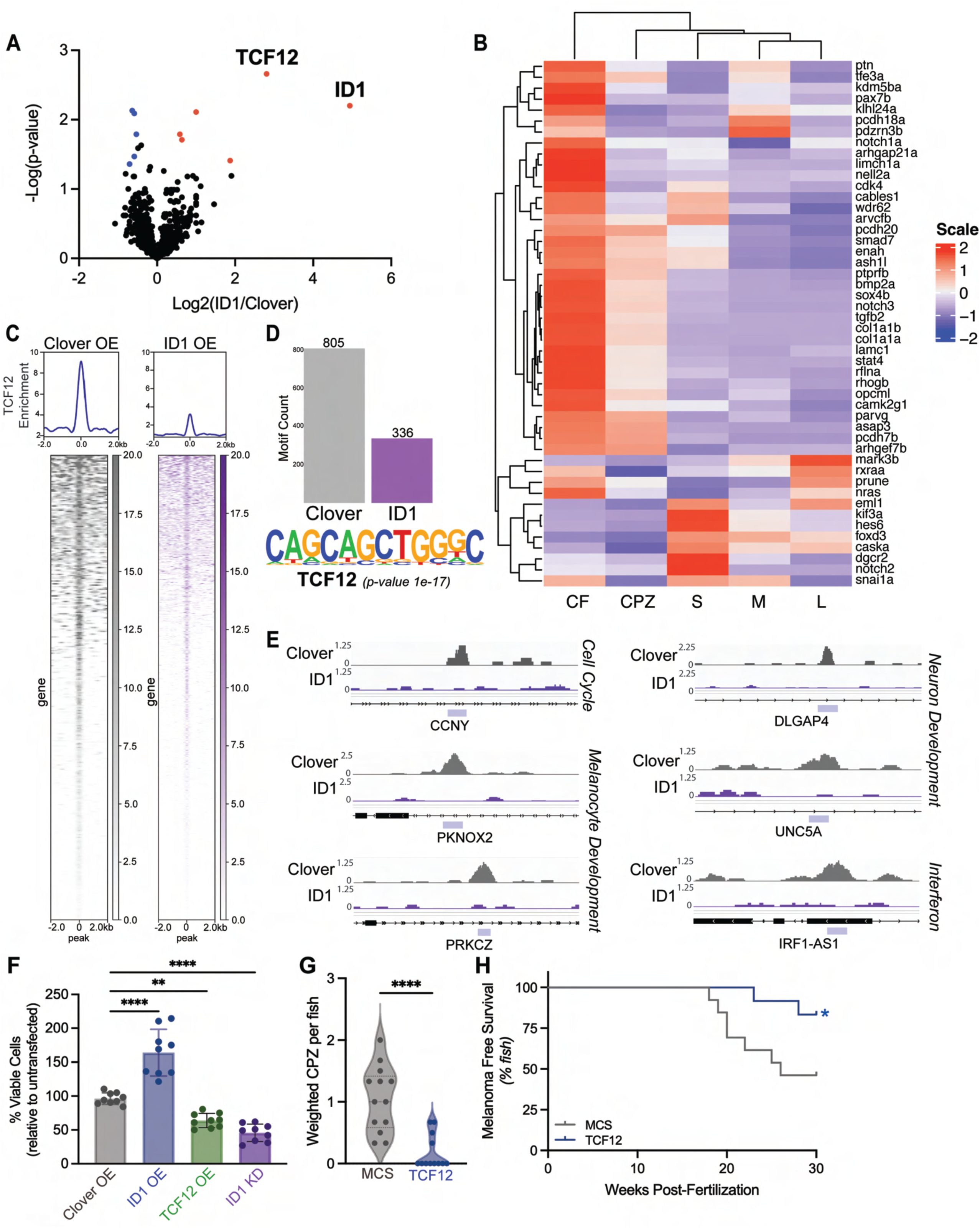
ID1 sequesters TCF12 from chromatin to drive cancer precursor zone formation through loss of development and signaling regulatory genes. **A**, Volcano plot analysis of IP-MS data comparing proteins bound to ID1 vs Clover pull down. **B**, Heatmap of TCF12 targets identified by their decreased expression in the context of ID1 overexpression and the presence of a TCF12 motif in the promoter/enhancer region. **C,** Heatmaps depicting ChIP-seq peak structure following Clover (left, gray) or ID1 (right, purple) overexpression in PMEL(BRAF^V600E^) cells. A kb window is centered on the peak center highlighting peaks lost after ID1 overexpression. **D,** Quantification of peaks with TCF12 motifs following Clover (n=805) or ID1 (n=336) overexpression. TCF12 motif is only enriched in Clover overexpressing cells (p = 1e-17). **E,** IGV tracks of TCF12 motif containing peaks in clover (gray) or ID1 (purple) overexpressing cells. Purple bar indicates peak called by MACS. **F,** Bar graph representing viable cells relative to non-transfected controls following Clover, ID1, or TCF12 overexpression or ID1 knockdown in PMEL(BRAF^V600E^) ***p≤0.01; ****p≤0.001. **G,** Violin plot showing the weighted number of cancer precursor zones following TCF12 overexpression relative to empty control. Each dot represents one fish; n=12. ****p≤0.001. **H,** Kaplan-Meier curve of melanoma incidence. Melanoma incidence is delayed in TCF12 overexpressing fish that develop CPZs. *p≤0.05, n=12.

To identify the repressed TCF12 genes, we next performed ChIP-seq following ID1 or clover overexpression in PMEL-BRAF melanocytes. ID1 overexpression led to a marked reduction in TCF12 chromatin occupancy and loss of TCF12 motif enrichment relative to control (Fig. 5C,D). This marked decrease focally affected melanocyte development and neuronal genes, including cell body, synapse regulation, and somatodendritic compartment regulation (Fig. 5E; Supp. Fig. 15E). Specifically, ID1-OE led to loss of TCF12 occupancy at known neuronal genes, and melanoma tumor suppressor genes, *DLGAP4* and *UNC5A*(*39-43*). In addition to known markers of the neural lineage, TCF12 occupancy was lost at established melanin production genes, such as *PKNOX2* and *PRKCZ*(*44*), (*45*), suggesting CPZs may represent unpigmented precursor lesions. During CPZ formation, melanocytes attain an aberrant morphology, including a loss of dendrites, which potentially achieved via a decrease in these targets.

To explore how modulation of the ID1/TCF12 axis alters cell viability, we measured proliferation in PMEL-BRAF cells. ID1 overexpression led to a significant increase in cell proliferation relative to control, likely due to the loss of development and cell cycle regulators (Fig. 5F). Conversely, TCF12 overexpression or siRNA knockdown of ID1 reduced cell viability.

Consistent with our findings in human cell lines, TCF12 overexpression in zebrafish using the MiniCoopR system led to a significant decrease in the number of CPZs formed and tumor onset compared to control (Fig. 5G,H). In both a zebrafish model and human immortalized melanocytes, modulation of TCF12 expression reduces proliferation and oncogenic potential. In total, these data support the hypothesis that ID1 binds to and inhibits TCF12, preventing it from activating downstream targets thereby enabling morphological and transcriptional changes to form oligoclonal cancer precursor zones, a novel initiating lesion in melanoma.

## Discussion

Our studies identify a cancer precursor zone (CPZ) as a spatially restricted, oligoclonal attractor state that precedes melanoma initiation. CPZs arise as contiguous fields of melanocytes undergoing coordinated clonal expansion, displaying altered morphology and distinct transcriptional profiles. These zones act as self-sustaining reservoirs of cells that generate transcriptional diversity, enabling individual clones to independently explore alternative cell-state trajectories. Rare, clone-restricted transitions within CPZs are marked by neural crest fate reactivation, after which a single clone dominates and initiates tumor formation. Importantly, these findings align with cell-of-origin studies showing transformation of interfollicular and differentiated melanocytes in the mouse tail(*13*).

CPZs are stabilized by BMP–ID1–TCF12 signaling, which suppresses terminal melanocyte differentiation and synchronizes transcriptional programs across neighboring cells, maintaining cells within a shared, stable transcriptional attractor. This signaling axis balances clonal fitness, enabling multiple clones to expand in parallel and persist as an oligoclonal population over extended periods. These findings are consistent with a model in which BMP signaling stabilizes adult stem and progenitor cells within a reversible “holding” state, preserving future plasticity by restricting premature fate commitment(*46-51*). Subtle differences in BMP–ID1 signaling or downstream transcriptional rewiring allow rare clones to escape the CPZ attractor, enter neural crest–like transcriptional trajectories, and acquire tumor-initiating potential, paralleling clonal hematopoiesis⁴³ and providing a mechanistic explanation for both long-term oligoclonality and the low frequency of malignant progression.

Zebrafish CPZ melanocytes closely resemble atypical melanocytic proliferative lesions in early-stage human samples, highlighting cross-species conservation and revealing a potential window for preventative therapy. CPZs function as dynamic reservoirs where developmental and transcriptional programs guide cell behavior rather than immediate tumor formation. Many CPZs remain arrested despite ongoing clonal expansion, but rare clones stochastically activate neural crest progenitor-like programs and transition toward tumorigenesis. These findings suggest that early interventions targeting CPZ dynamics could suppress tumor initiation before malignant transformation occurs.

More broadly, our results support the cancer attractor state model, showing that tumor initiation is governed by entry into, maintenance of, and rare escape from a defined precursor state. Here, we show this precursor state is clonally derived from mature tissue populations. Oligoclonal precursor fields, like CPZs, have also been described in Barrett’s esophagus, ductal carcinoma in situ, prostatic intraepithelial neoplasia, the uterine cervix, and familial adenomatous polyposis(*52-56*). Together, these observations indicate that malignancy often emerges from stable multicellular precursor states that channel evolutionary trajectories over time. By resolving CPZ dynamics in vivo, our work reframes melanoma initiation as a multistep process shaped by attractor stability and clone-specific escape events. This model highlight opportunities for early intervention by stabilizing precursor states or preventing escape, thereby providing a framework for cancer prevention strategies targeting transcriptional and epigenetic states.

## Author Contributions

A.M.M., M.H.C., and L.I.Z. conceived of the study; L.I.Z. and G.F.M. supervised the research; A.M.M. and M.H.C. designed and performed experiments with the assistance of H.R.N., J.B., E.W., I.F.G., E.L., K.D.D., C.S.B., and P.A.; J.K.M. assisted with histology and pathologic assessment of fish and human samples; A.M.P., S.Y., C.L. performed computational analysis for all sequencing experiments; C.G.L. A.E., M.D., M.S.S., S.D., J.C., I.W., provided human samples and pathologic analysis. M.G., Y.K., K.J.L performed immunofluorescent staining and analysis on human samples.

## Acknowledgements

We would like to thank Christian Lawrence, Shane Hurley, Li-Kun Zhang, Kara Maloney, and Hannah Nations for expert fish care. The authors thank many of their colleagues for their discussions and critical reading of the manuscript. We thank Dana-Farber/Harvard Cancer Center for the use of the Specialized Histopathology Core, which provided histology and immunohistochemistry service. The authors would like to thank the Neurobiology Imaging Facility (NIF) at Harvard Medical School for performing the RNAscope and the Single Cell Biology core for assisting with the sequencing of the single cell atlas. We further acknowledge the assistance the Translational Research Institute Microscopy Core Facility at the University of Queensland. The Dana-Farber/Harvard Cancer Center is supported in part by an NCI Cancer Center Support Grant #P30CA06516. We thank Ross Tomaino and the Harvard Medical School Taplin Mass Spectrometry Facility for their assistance in performing and analyzing our mass spectrometry experiments. This work was supported by R01CA103846-14, American Cancer Society Postdoctoral Fellowship PF-18-150-01-DCC (A.M.M.), K00CA253749 (M.H.C.), and F32CA294934 (K.D.D.). L.I.Z. is a Howard Hughes Medical Institute Investigator. This work was also supported by Lyon Integrated Research Institute in Cancer (J.C., S.D.), Institut Convergence

