## Supplemental Info for "BMP-dependent oligoclonal cancer attractor state precedes neural crest fate in melanoma initiation"

#### SUPPLEMENTARY INFORMATION

##### Materials and Methods

*Generation of BRAF;p53 zebrafish*

*mitfa:BRAF*$^{V600E}$*;p53*$^{-/-}$*;mitfa*$^{-/-}$*;crestin:EGFP;mcr:Empty;mitfa:mCherry;tyr*$^{-/-}$ zebrafish (referred to as *BRAF;p53*) were generated by injecting *mcr:Empty* and *mitfa:mCherry* at 25 ng/μL, 5 mg/mL Cas9 protein (PNA Bio CP02), a gRNA targeting *tyr* (GGACTGGAGGACTTCTGGGG) at 50 ng/μL, and Tol2 mRNA at 20 ng/μl into the single cell stage of *mitfa:BRAF*$^{V600E}$*;p53*$^{-/-}$*;mitfa*$^{-/-}$*;crestin:EGFP* embryos(*57*). These fish were originally developed to be used in combination with MiniCoopR (mcr), which is a transgenic tool that permits selective mis-expression of any gene in melanocytes, thereby allowing us to assess the ability of each gene to promote or inhibit melanoma initiation(*18*). Injection of the MiniCoopR vector mosaically rescues melanocyte development via an *mitfa* minigene, allowing melanomas to form. The MiniCoopR vector also contains the open reading frame of any candidate gene driven specifically in melanocytes by the *mitfa* promoter, which enables us to assess its oncogenic potential. For the morphological and molecular characterization experiments, a MiniCoopR vector containing only the *mitfa* minigene was injected to rescue melanocyte formation (MCR:MCS).

*mitfa:BRAF*$^{V600E}$*;p53*$^{-/-}$*;mitfa*$^{-/-}$*;crestin:EGFP* zebrafish were generated by crossing *mitfa:BRAF*$^{V600E}$*;p53*$^{-/-}$*;crestin:EGFP*[3] zebrafish with *mitfa:BRAF*$^{V600E}$*;p53*$^{-/-}$*;mitfa*$^{-/-}$ zebrafish. *mitfa:mCherry* was cloned by Gateway reaction using the zebrafish *mitfa* promoter and the Tol2 mCherry middle entry plasmids. The *tyrosinase* gRNA was synthesized using SP6 in vitro transcription(*19, 58*). In brief, the *tyr* oligo template

(CCTCCATACGATTTAGGTGACACT ATAGGACTGGAGGACTTCTGGGGTTTTAGAGCTAGAAATAGCAAG) and the constant oligonucleotide (AAAAGCACCGACTCGGTGCCACTTTTTCAAGTTGATAACGGACTAGC CTTATTTTAACTTGCTATTTCTAGCTCTAAAAC) were annealed and filled in with T4 DNA polymerase (New England BioLabs, M0203S). The product was PCR amplified, gel purified and transcribed using MEGAscript T7/SP6 (ThermoFisher Scientific, AM1333). gRNAs were cleaned up using Direct-zol RNA Miniprep kit (Zymo Research, R2051).

To generate the *casper;mcr:NRAS*$^{Q61R}$ or *casper;mcr:BRAF*$^{V600E}$ zebrafish, *mcr:NRAS*$^{Q61R}$ or *mcr:BRAF*$^{V600E}$ was injected into *casper;crestin:EGFP* embryos along with Tol2 mRNA and *mitfa:mCherry* at a concentration of 25 ng/µL.

This study was performed in strict accordance with the recommendations in the Guide for the Care and Use of Laboratory Animals of the National Institutes of Health. The animal research protocol was approved by the Institutional Animal Care and Use Committee of Boston Children's Hospital. All zebrafish used in this study were maintained and euthanized under the guidelines of the Institutional Animal Care and Use Committee of Boston Children's Hospital.

*Histology*

Fish were euthanized and fixed in 4% paraformaldehyde overnight at 4°C. Paraffin embedding, sectioning, Hematoxylin and Eosin (H&E) staining were performed according to standard techniques by the Brigham & Women's Hospital Pathology Core. Immunohistochemistry was performed with 5 µm thick formalin-fixed, paraffin-embedded tissue sections using the Leica Bond III automated staining platform and the Leica Biosystems Refine Detection Kit. Antibody mCherry (Thermo Fisher #M11217) was run at 1:200 dilution with EDTA antigen retrieval and Goat Anti-Rat IgG secondary antibody from Vector Labs catalog number PI-9401-.5. Antibody GFP from Abcam, catalog number 6556, polyclonal, was run at 1:800 dilution with EDTA antigen retrieval. Antibody PCNA (CST #2586), was run at 1:16000 dilution with citrate antigen retrieval. Antibody Phospho ERK (p44/42 MAPK; CST #4370), was run at 1:150 dilution with citrate antigen retrieval. All human tissue samples were derived from the Pathology Archives of the Brigham and Women's Hospital, University of San Francisco, The University of Queensland, or Lyon Sud Hospital with full institutional review board approval. Patient consent for experiments was not required because de-identified pathological specimens of human samples are discarded material by our institution, and thus the studies were exempt. Immunofluorescence studies were performed on paraffin-embedded sections of formalin-fixed tissue. Tissue sections cut at 5 µm intervals were deparaffinized, rehydrated, and heated with Target Antigen Retrieval Solution (Dako, Agilent Technologies) in a pressure cooker. Sections were blocked with 10% animal serum for 30 min before incubation with primary antibodies. Samples were incubated overnight at 4°C with the following primary antibodies: mouse IgG2a anti-ID1 (1:500; SCBT, sc133104), mouse IgG2b anti-MART1 (1:1, Biolegend #917902). Sections were then treated with 0.1% Sudan Black (Abcam) for 10 minutes to remove autofluorescence. The following secondary antibodies were used: goat anti-mouse IgG2b (1:2000; Invitrogen #A-21141), goat anti-mouse IgG2a (1:2000; Invitrogen #A-21135), goat anti-mouse IgG1 (1:2000; Invitrogen #A-21235). Slides were mounted with Floromount G with NucBlue. For samples in Figure 4C, the sections underwent immunofluorescence staining using the OPAL™ technology (Akoya Biosciences) on a Leica Bond RX with DAPI (Akoya OPAL kit), SOX10 (SCBT sc-365692, 1:1000), MITF (Sigma, 284M-96, 1:200), or ID1 (Cliniscience sc-133104, 1:1000). Sections were digitalized with a PhenoImager HT scanner (Perkin Elmer). ID1 expression in melanocytic cells was evaluated using the HALO AI™ Software (Indica Labs), with HighPlex FL module. Briefly, a threshold for ID1 expression has been settled to define negative, low and high expression, and two cell phenotypes were characterized as melanocyte (DAPI+, MITF+ and/or SOX10+) or ID1+ melanocyte (DAPI+, MITF+ and/or SOX10+, ID1+).

Whole transcriptomic spatial profiling of human melanocytic lesions was performed using the 10x Genomics Visium CytAssist platform as previously described(*59, 60*). Since each 'spot' may contain several cells depending on cell size, spot deconvolution was performed using a single-cell skin whole transcriptomic dataset generated following the 10x Genomics Chromium X workflow. Spots enriched for melanocytes were assessed for *ID1* and *ID2* expression using a module score analysis using the Seurat v5 R package, and the *AddModuleScore*. Spots were grouped based upon their histopathological diagnosis (Naevus, MIS, and TM). Summary statistics including the mean, standard deviation (SD), and median expression per diagnosis group were calculated using R (dplyr package). The module score represents a relative expression value of *ID1* / *ID2* in each diagnostic category, which provides a normalized measure of gene expression level that accounts for background expression variability. A representative image demonstrating *ID1* / *ID2* gene expression within the spots enriched with melanocytes was visualized using *SpatialFeaturePlot*.

*In Situ Hybridization (RNAscope)*

RNAscope Multiplex Fluorescent Assay(*33*) (Biotechne) was performed on formalin-fixed paraffin embedded cancerized field, cancer precursor zone, patch, and tumor sections. Melanoma stages were assigned by H&E and mCherry or GFP expression (IHC methods described above). Protocol was followed according to manufacturer's instructions except protease 3 was used instead of protease 4. Each sample was hybridized with zebrafish id1 (#517531, C1 probe, red) and zebrafish crestin (#534061, C2 probe, far red) alongside a DAPI stain to mark nuclei. RNAscope was provided by the Neurobiology Imaging Facility (NIF) at Harvard Medical School. Stained slides were imaged with a 40x objective on a Nikon Eclipse Ti-2 spinning disk confocal microscope. All images were acquired using NIS-Elements (Nikon) and minimally processed using Imaris.

*Imaging*

Zebrafish were anesthetized with 4% MS-222 (Pentair, TRS1) and imaged on a Nikon SMZ18 Stereomicroscope or a Nikon C2si Laser Scanning Confocal using a 10x objective. Histologic sections were imaged on a Nikon Eclipse Ti-2 Spinning Disk Confocal using a 40x or 100x objective. Maximum intensity projections of Z stacks or three-dimensional re-constructions are presented here. Images were minimally processed using Photoshop, FIJI, or Imaris. Multiple tiled images of adult zebrafish and histology were stitched together by using the automated Photomerge function in

Photoshop. Given the variability in the rescue specific to each animal, number of zones/patches per fish were weighted for *mitfa:mCherry* rescue.

*FACS Cell Isolation*

*mitfa:BRAF*$^{V600E}$*;p53*$^{-/-}$*;mitfa*$^{-/-}$*;crestin:EGFP;mcr:Empty;mitfa:mCherry;tyr*$^{-/-}$ zebrafish from the same cohort were categorized by stage: control cancerized field, cancer precursor zone, small *crestin* patch, medium *crestin* patch, and large *crestin* patch/tumor. The zones and patches were manually dissected and individually chopped for 1-2 minutes. The finely chopped tissue was suspended in 3 mLs of TrypLE Express (ThermoFisher Scientific, 12605028) and incubated at 37$^{o}$C shaking at 300 rpm for 30 minutes. Samples were filtered through a 40 µm filter and washed with 5 mL of FACS buffer, consisting of DPBS (ThermoFisher Scientific, 14190144), 10x Penicillin Streptomycin (ThermoFisher Scientific, 15140122), and 2% heat-inactivated FBS (ThermoFisher Scientific, A3840001). Samples were centrifuged at 500 rcf for 5 minutes and resuspended in 150 µL FACS buffer. Samples were stained with SYTOX blue (ThermoFisher, S34857) or DRAQ7 (Abcam, ab109202) immediately before sorting for live (dye negative), mCherry and/or EGFP positive cells on a FACSAria II (BD Biosciences). FACS cell isolation was used for bulk and single cell RNA-seq, ATAC-seq, GESTALT barcode, and transplantation experiments.

*GESTALT Barcoding and Analysis*

GESTALT barcoding and downstream analyses were performed as previously described(*21, 22*). Single cell embryos resulting from the cross of a GESTALT barcode female and GESTALT guide male were injected with MCR:BRAF, mitfa:mCherry, p53 and albino gRNA to generate mosaic $BRAF^{V600E}$;$p53^{-/-}$ $mCherry^{+}$ melanocytes along with barcode sites 1-4 gRNA to induce editing of the first 4 GESTALT sites. For BMP studies, MCR:MCS (control), MCR:ID1, MCR:caSMAD1(DVD), or MCR:dnBMPR were injected along with the oncogenic components and sgRNA for sites 1-4 as described above. At 48 hpf, embryos were heated for 45 min at 37°C to induce editing of GESTALT sites 5-9. Barcoded fish were monitored for induction of CPZ and tumor formation using the imaging parameters described above. Melanoma initiation stages were isolated via FACS, as detailed above, and efficiency editing was confirmed by T7 endonuclease I assay and CRISPR-targeted sequencing of the GESTALT barcode. Read mapping, insertion/deletion calling, and downstream barcode analysis was performed as described(*22, 61*). Barcodes with less than 10 mapped reads or detected in more than 25% of fish were discarded from analysis. Dominant clones are at least three standard deviations larger than the mean size of the cancerized field control (>20% of reads), a similar cutoff as described in a zebrafish model of hematopoesis(*62*).

*scRNA-seq*

For the 10X platform, cancerized field (n=2), cancer precursor zones (n=4), and crestin patches (n=2) were dissociated into a single cell suspension and FACS sorted for viability. Viable cells were GEM barcoded using the Single Cell 3’ Reagent Kit to target 10,000 cells per sample. cDNA was generated and the library was constructed following the 10X protocol. Libraries were sequenced on an Illumina NovaSeq by the Single Cell Core at Harvard Medical School. For InDrop, cells isolated from the skin of 6 control fish and 4 fish with small *crestin* patches were FACS sorted for *mitfa:mCherry*, as described above. 1360 melanocytes were combined with 29,000 mCherry negative skin cells in the control and 6224 melanocytes were combined with 24,000 mCherry negative skin cells in the *crestin* patch sample to yield 2 samples each with 30,000 cells. These samples were run through the inDrops platform by the Single Cell Core at Harvard Medical School, as previously described(*63, 64*). Libraries were run on an Illumina HiSeq 4000 with paired end 150bp. inDrops single-cell RNA-seq analysis follows the instruction as described in https://github.com/indrops/indrops. For both platforms, FASTQ files were processed and aligned to GRCz11 genome. Data were normalized and processed using the Seurat and monocle3 packages in R(*65-67*).

*RNA-seq*

RNA was extracted from 5,000 sorted melanocytes from 3-4 control or ID1-overexpressing fish per stage. Ultralow input RNA-seq was performed using the SMART-Seq v4 Ultra Low Input RNA kit for Sequencing (Clontech, 634888) and Nextera XT DNA Library Preparation Guide (Illumina, FC-131-1024). Libraries were run on an Illumina HiSeq 4000 with paired end 150bp. Quality control of RNA-Seq datasets was performed by FastQC and Cutadapt to remove adaptor sequences and low-quality regions(*68, 69*). The high-quality reads were aligned to Ensembl build GRCz11 of zebrafish genome using Tophat 2.0.11 without novel splicing form calls(*70*). Transcript abundance and differential expression were calculated with Cufflinks 2.2.1(*71*). Differential expression analysis was performed using DESeq2 in R[46]. Pathway analysis was performed using Metascape and graphed using GraphPad Prism 7 (GraphPad Software)(*72*).

PMEL(hTERT/CDK4(R24C)/p53DD)) with and without $BRAF^{V600E}$ cells (courtesy of D. Fisher lab(*34*)) were harvested at 90% confluency in trizol from three biological replicates. RNA was obtained using Zymo Direct-Zol RNA Miniprep Kit and integrity determined using Agilent RNA Screentape (RIN >9). 500 ng of RNA was used as input for rRNA depletion with the NEBNext rRNA Depletion Kit v2 (human/mouse/rat) (NEB E7400). Samples were then converted to cDNA libraries using NEBNext Ultra II RNA Library Prep Kit for Illumina (NEB E7770). Libraries were sequenced at 360M paired-end, 2x150bp reads via Azenta Life Sciences. The standard alignment and Deseq2 pipelines were run as described above.

*ATAC-seq*

5,000 sorted melanocytes from 3-4 control or ID1-overexpressing fish at each stage were lysed and subjected to “tagmentation” reaction and library construction as previously described(*73*). Libraries were run on an Illumina HiSeq 4000 with paired end 150bp. All zebrafish ATAC-Seq datasets were aligned to build version Ensembl build GRCz11of the zebrafish genome using Bowtie2 (version 2.2.1) with the following parameters: --end-to-end, -N0, -L20(*74*). We used the MACS2 (version 2.1.0) peak finding algorithm to identify regions of ATAC-Seq peaks and derive the normalized tracks, with the following parameter --nomodel --shift -100 --extsize 200 --SPMR(*75*). A q-value threshold of enrichment of 0.05 was used for all datasets. The motifs enriched in ATAC-seq peaks of interest were analyzed using the findMotifsGenome program in the HOMER package(*76*). The corresponding zebrafish genome sequences are used as background sequences in motif search. The top known HOMER motifs and de novo motifs with q-value less than *** are calculated. HOMER motif results are presented as a heatmap of the percent of targets containing the indicated motif minus background. For example, 41.8% of new or increased target peaks in large *crestin*+ samples contain a RUNX motif, as compared to 20.7% of background peaks. Therefore, the heatmap indicates 21.1% of target peaks are enriched for a RUNX motif.

*Transplantation*

FACS isolated *mitfa:mCherry*+/*crestin:EGFP*- cancer precursor zone cells or *mitfa:mCherry*+/*crestin:EGFP*+ cells from small *crestin* patches or tumors were enrobed in 3 µL Matrigel (Corning, 356234) and injected at concentrations of 5,000, 3,000, 1,000, or 500 cells under the skin of *casper* recipient fish irradiated sub-lethally with 30 Gy split over 2 days. Isolated cells from each donor went into 3-7 recipient fish. Fish were imaged as described above and limit dilution calculations and curves were generated using the Extreme Limiting Dilution Analysis software (http://bioinf.wehi.edu.au/software/elda/)[49] on data collected 14 days post-transplant.

*IP-MS*

The fusion (no stop) human ID1 or Clover open reading frames were cloned into the pcDNA3.2 V5 destination vector (Invitrogen, 12489019) to create V5 tagged ID1 or Clover constructs. Constructs were transiently transfected into A375 human melanoma cells using Lipofectamine®3000 (Invitrogen, L3000001) in 10 $cm^2$ plates with three independent replicates. 48 hours after transfection, cytoplasmic and nuclear fractions were isolated using the NE-PER™ Nuclear and Cytoplasmic Extraction Kit (ThermoFisher, 78833) and lysed per protocol. Anti-V5 (Clone V5-10, Sigma, V8012) was conjugated to protein G beads (ThermoFisher, 10004D). IP'd proteins were eluted and submitted for mass spectrometry using the Taplin Mass Spectrometry Facility at Harvard University. Proteins were included in analysis if all replicates had greater than 3 peptides.

*ChIP-Seq*

PMEL(hTERT/CDK4(R24C)/p53DD)/BRAF(V600E) cells (courtesy of D. Fisher lab(*34*)) were transiently transfected with V5-Clover or V5-ID1, as detailed above, in 15 $cm^2$ plates with three independent replicates. Cells were crosslinked in 11% formaldehyde, lysed, and sheared as described(*77*). Solubilized chromatin was immunoprecipitated with 10 µg TCF12 antibody (SCBT, sc357). Antibody-chromatin complexes were purified and libraries were prepared as described(*77*). All ChIP-Seq datasets were aligned to Ensembl build version GRCh38 of the human genome using Bowtie2 (version 2.2.1) with the following parameters: --end-to-end, -N0, -L2086(*74*). MACS2 (version 2.1.0) peak finding algorithm was used to identify regions of ChIP-Seq peaks, with a q-value threshold of enrichment of 0.05 for all datasets(*75*). ChipSeeker is utilized to annotate ChIP-seq peaks to neighboring genes according to Ensembl gene annotation(*78*). The parameters are defined as proximal promoter: 500 bp upstream – 50 bp downstream of TSS; distal promoter: 2kb upstream – 500 bp downstream of TSS; enhancer: 100 kb from TSS. The genome-wide occupancy profile figures were generated by deeptools2 with peaks centered(*79*). HOMER analysis was performed to confirm transcription factor binding under peaks with hg38 genome utilized as a random set of background peaks for motif enrichment(*76*). To count TCF12 motifs enriched in called peaks, mapped TCF12 motifs across the hg38 genome from ENCODE  enrichment was used(*80*). Reactome pathway analysis was completed with Gene Ontology(*81, 82*).

*Cell Viability*

The fusion human TCF12 open reading frame was cloned into pcDNA3.2 V5 destination vector to create V5 tagged TCF12. A375 or PMEL cells were seeded in opaque-walled 96-well plates. 5,000 cells were transiently transfected with V5-ID1, V5-TCF12 or a V5-clover control for 48 h. For siRNA knockdown, RNAiMax lipofectamine was added to siRNA targeting ID1, allowed to complex, and added to 5,000 A375 cells for 48 h. Cell viability was then measured using CellTiter-Glo (Promega #G7570) and luminescence was read on Synergy Neo plate reader. Experiments were performed in technical triplicate for each biological replicate (n=3).

*Statistics*

A two-way ANOVA with Tukey's multiple comparisons test was used to compare the weighted number of cancer precursor zones or *crestin* patches and the immunofluorescence quantitation. The calculations were performed using GraphPad Prism 7 (GraphPad Software).

#### Supplemental Figure 1

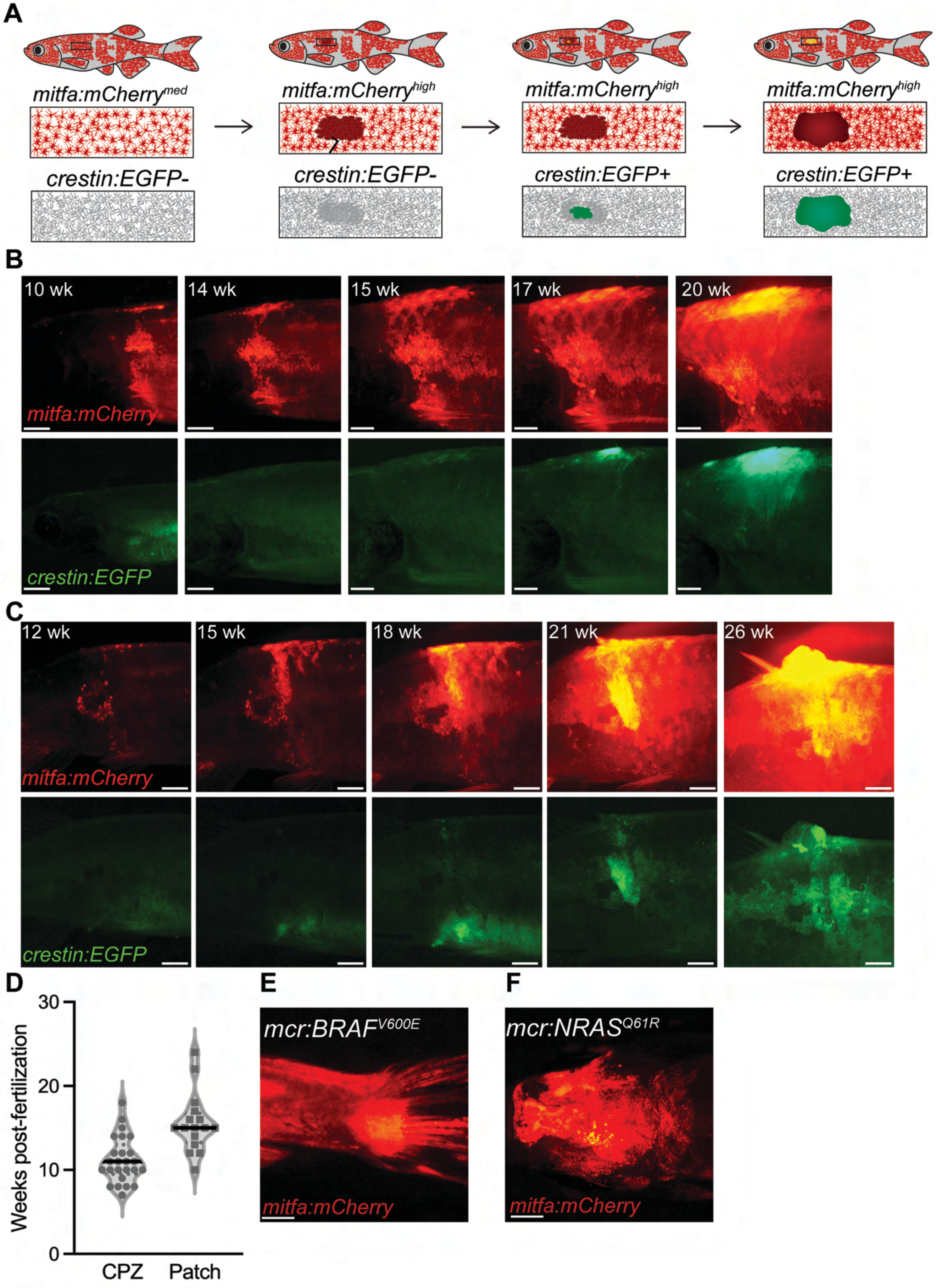

**Supplemental Figure 1 | Cancer precursor zone formation precedes neural crest reactivation in zebrafish. A**, Melanoma initiates in a stage-specific manner, beginning with the formation of an *mitfa*$^{high}$ *crestin:EGFP*- cancer precursor zone. The neural crest progenitor state is reactivated from within the cancer precursor zone and tumor formation occurs following this reactivation. **B,** Melanoma initiation stages in a zebrafish with a dorsal tumor. Scale bar: 1mm**. C** , Images of melanoma initiation for zebrafish marked * in Fig. 1A. Scale bar: 1mm. **D,** Violin plot of CPZ and patch onset. CPZ have a median onset of 11 wpf (circle, left) while patch occur at 15 wpf (square, right). Median is shown in solid black line. Each dot is a fish; n=25. **E,F,** Cancer precursor zones form in *mcr:BRAF*$^{V600E}$ (**E**) and *mcr:NRAS*$^{Q61R}$ (**F**).

**Supplemental Figure 2**

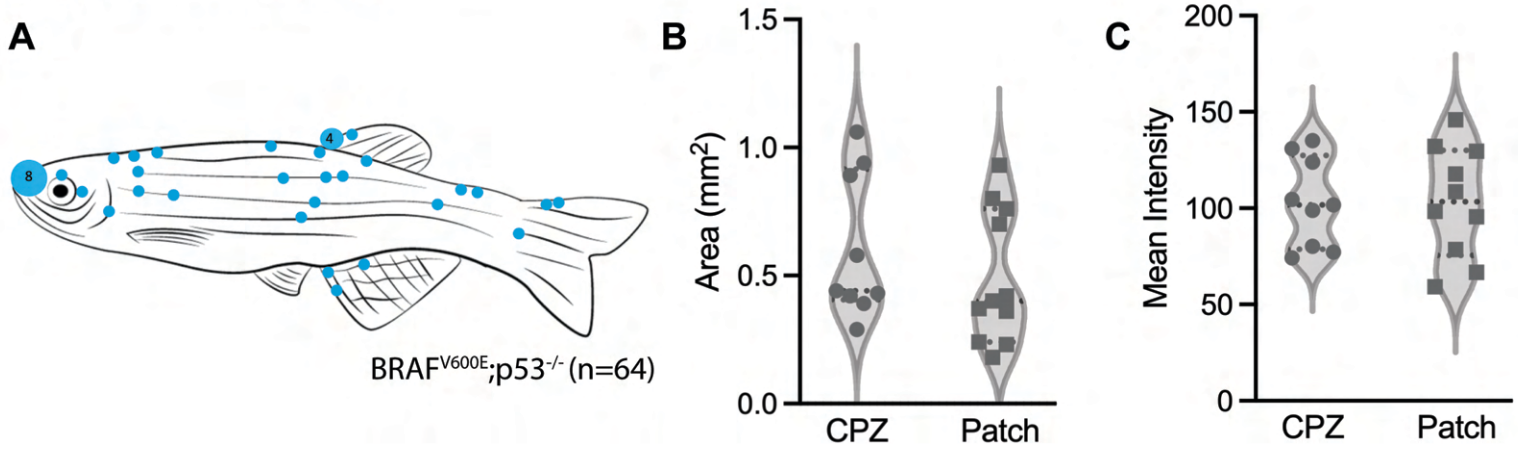

**Supplemental Figure 2 | Neural crest reactivation is not dependent on anatomic position, size, or *mitfa:mCherry* intensity. A,** CPZ anatomic location across the zebrafish body. CPZ incidence does not correlate with anatomic location. Size of the dot represents the count of CPZs at that location with the smallest dot being n=2, n=64. **B,C.** Violin plots of cancer precursor zone area (**B**) or mean mitfa:mCherry intensity within that area (**C**). Size or mitfa:mCherry intensity does not predict which CPZs will reactivate neural crest (square; mean area: 103.2±30.0; mean intensity: 0.49±0.26 ) compared to CPZs that stall in the CPZ stage (circle; mean area: 0.60±0.28; mean intensity: 103.0±23.1). Each point is a fish, n=20.

#### Supplemental Figure 3

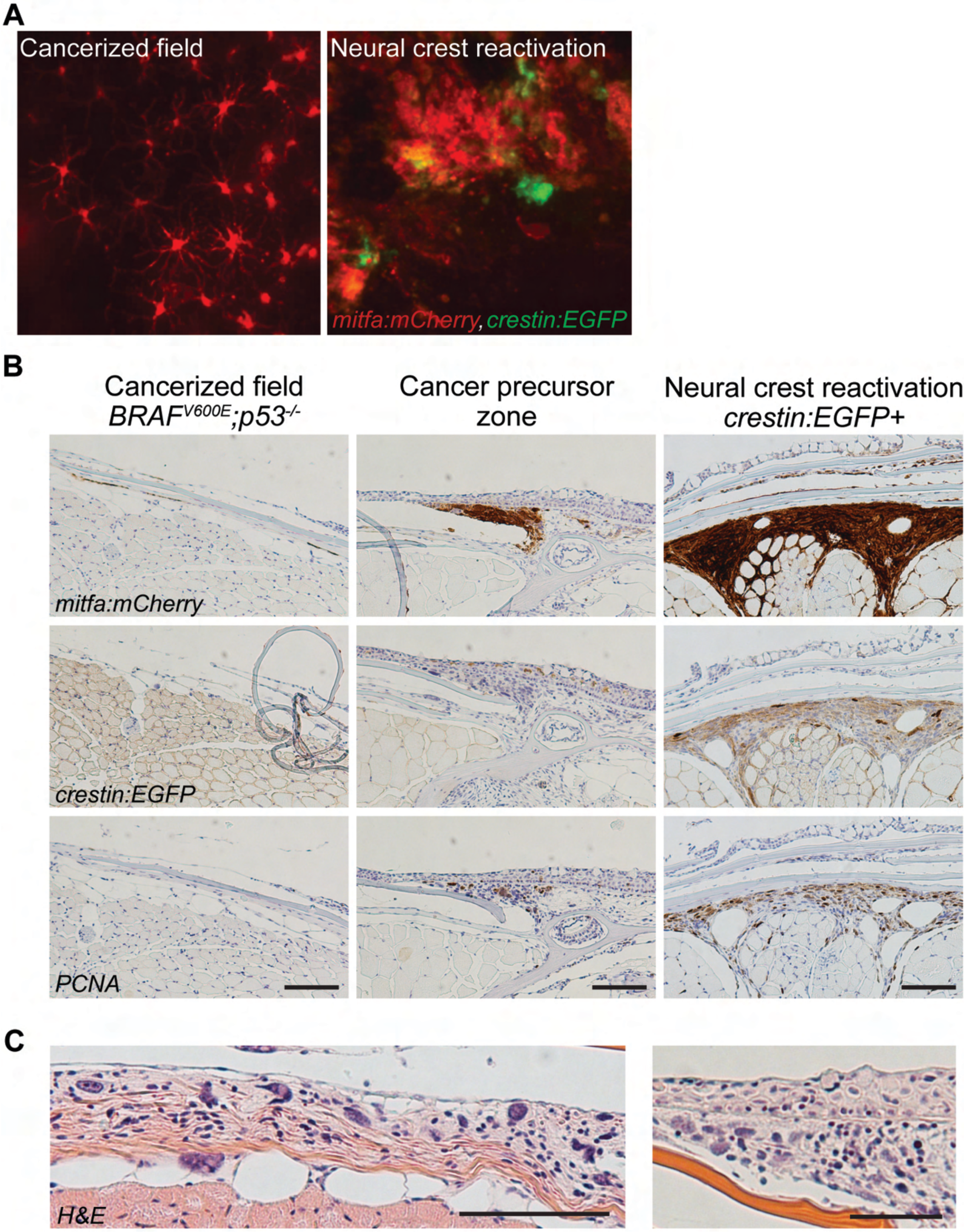

**Supplemental Figure 3 | Cancer precursor zones are proliferative regions of atypical melanocytes that precede neural crest reactivation. A**, Confocal images depicting morphological changes; cancerized field melanocytes are dendritic and regularly spaced, whereas patches with neural crest reactivation are made of atypical non-dendritic melanocytes. **B**, PCNA staining reveals that cancer precursor zones and patches with neural crest reactivation are highly proliferative compared to infrequently proliferative cancerized field melanocytes. Scale bars, 100 μm. **C**, H&E staining of cancer precursor zones show that nuclei are dysplastic. Scale bars, 100μm.

#### Supplemental Figure 4

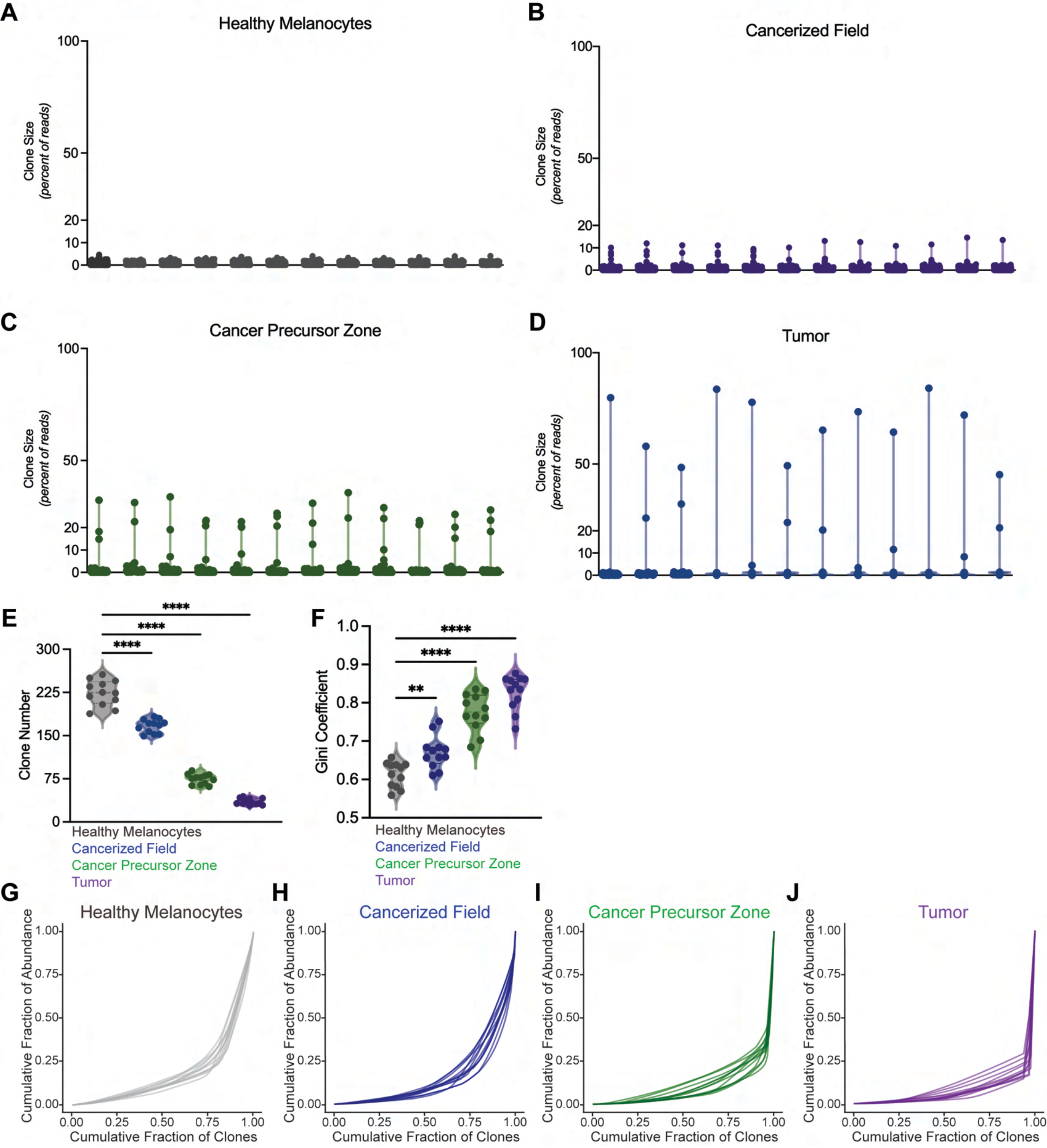

**Supplemental Figure 4 | Clonal dynamics of melanoma initiation stages. A-D,** Violin plot of clone size (calculated by percent of reads) from individual GESTALT barcoded fish with healthy skin (**A**), cancerized field (**B**), CPZ (**C**), and tumor (**D**) melanocytes from Fig. 1D-F. Each dot is a clone and each line is a fish. **E, F**, Violin plot of total clone number (**E**) and Gini coefficient (**F**) in a cohort of GESTALT barcoded fish across melanoma initiation stages (n=12 fish). **, $p \leq 0.01$; ****, $p \leq 0.0001$. **G-J**, Lorenz curves of cumulative fraction of abundance and fraction of clones for healthy (**G**), cancerized field (**H**), CPZ (**I**), and tumor (**J**) melanocytes.

#### Supplemental Figure 5

**A**

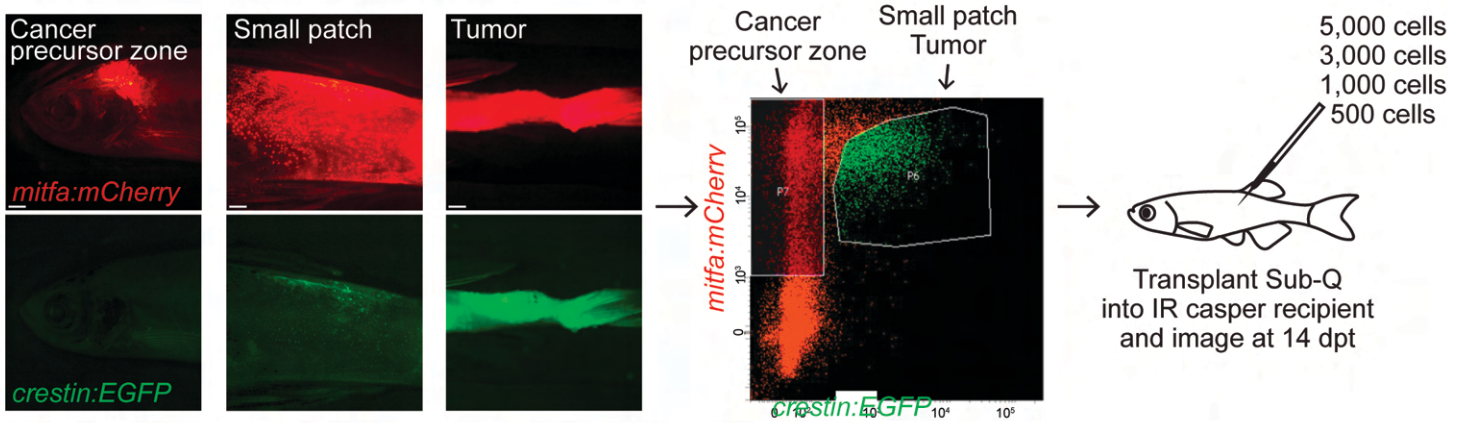


**B**

| **Group** | **Lower** | **Estimate** | **Upper** |
|---|---|---|---|
| mCherry Skin | 99458 | 13345 | 1818 |
| Cancerized Field | 4090 | 1236 | 374 |
| Cancer Precursor Zone | 1126 | 806 | 578 |
| Patch | 881 | 373 | 158 |
| Tumor | 564 | 348 | 217 |

**Supplemental Figure 5 | Malignant potential of melanoma initiation stages. A**, Schematic of experiment. Precursor lesion (*mitfa:mCherry+/crestin-*), small *crestin* patch (*mitfa:mCherry+/crestin+*), or tumor cells (*mitfa:mCherry+/crestin+*) were isolated via FACS and subcutaneously transplanted into irradiated (IR) recipients at various densities. **B**, Confidence interval table for 1/(engraftment cell frequency) in transplant studies.

### Supplemental Figure 6

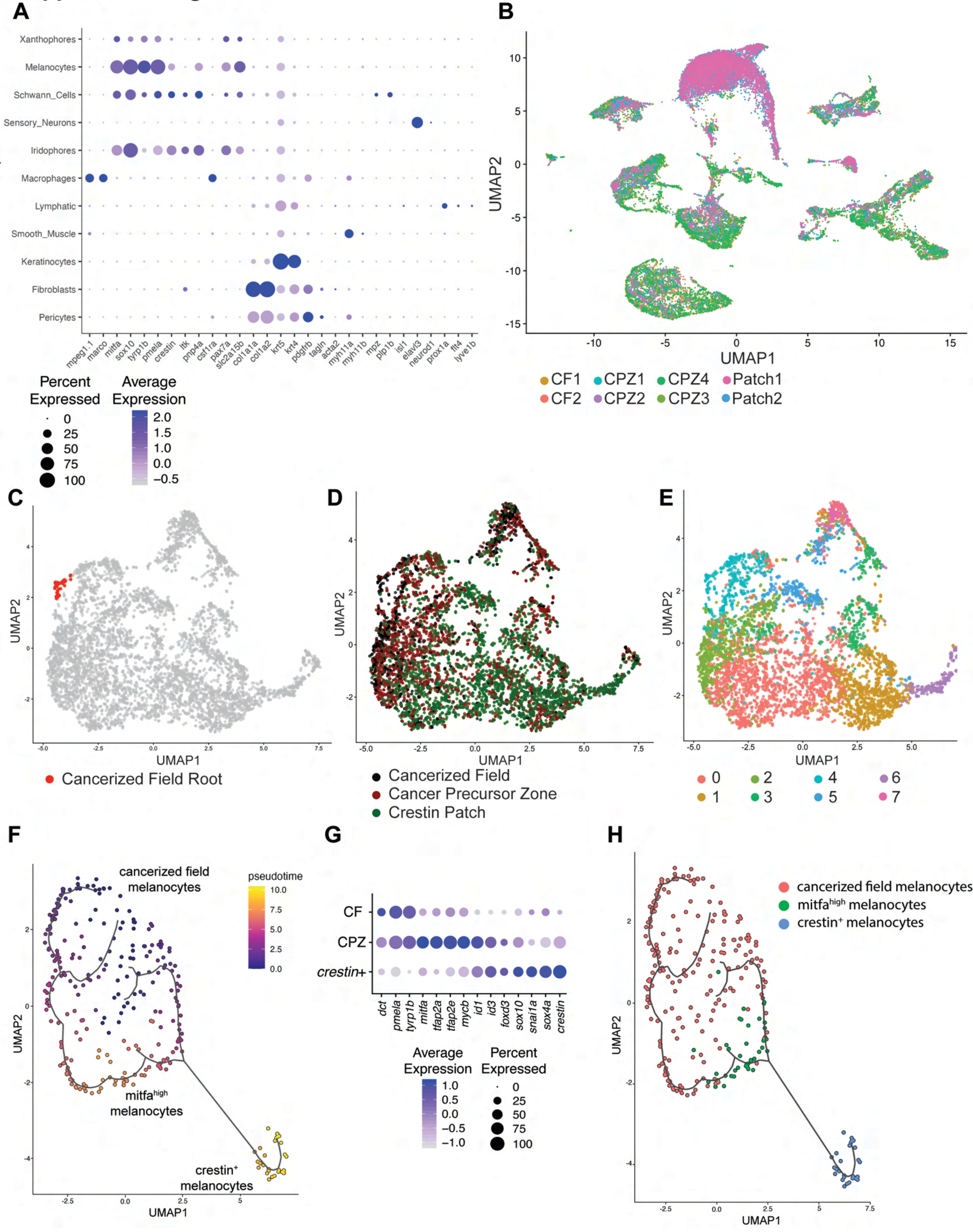

**Supplemental Figure 6 | Single cell transcriptome of melanoma initiation. A**, Dot plot showing the expression of cell-type specific genes used to identify clusters. **B**, UMAP visualization of cell clusters marked by sample. The control sample contains cells isolated from two cancerized field fish, four CPZ fish, and 2 crestin patch fish. **C,** UMAP of melanocytes (gray) with the highlighted cancerized field root (red) used to calculate pseudotime. **D,** UMAP of the assigned cell identities from the Pseudotime presented in Fig. 2G-I. **E,** UMAP of the assigned Seurat transcriptional clusters. **F**, UMAP of pseudotime analysis from InDrop data with cancerized field melanocytes as the root. **G**, Dot plot of differential genes expressed in cancerized field control melanocytes (CF), cancer precursor zone (CPZ), and crestin+ melanoma subsets. **H**, UMAP of the assigned cell identities from (**F**).

### Supplemental Figure 7

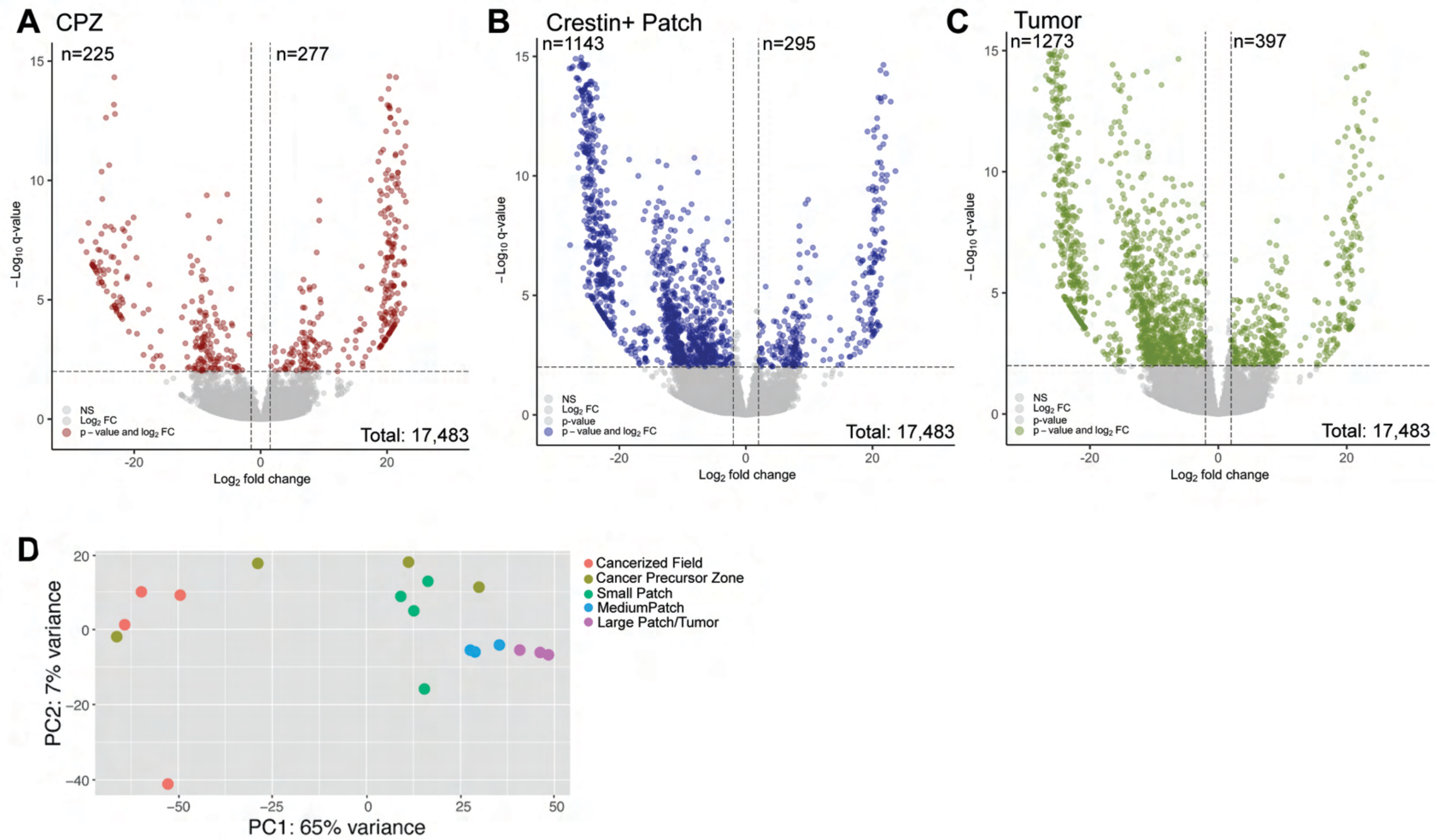

**Supplemental Figure 7 | Differential gene expression across melanoma stages. A-C**, Volcano plots showing differentially expressed genes in CPZ (**A**), crestin patch (**B**), and tumor (**C**) relative to cancerized field control. Dashed lines represent a 2 $log_2$FC (over cancerized field) and a 10e-3 q-value as assigned by DeSeq2. Colored regions indicate genes that meet the logFC and q-value threshold. **D**, PCA plot of cancerized field, cancer precursor zones, and patches. CPZ melanocytes (yellow) represent a transitory population from cancerized field (red) to patch (green and blue) and tumor (purple).

#### Supplemental Figure 8

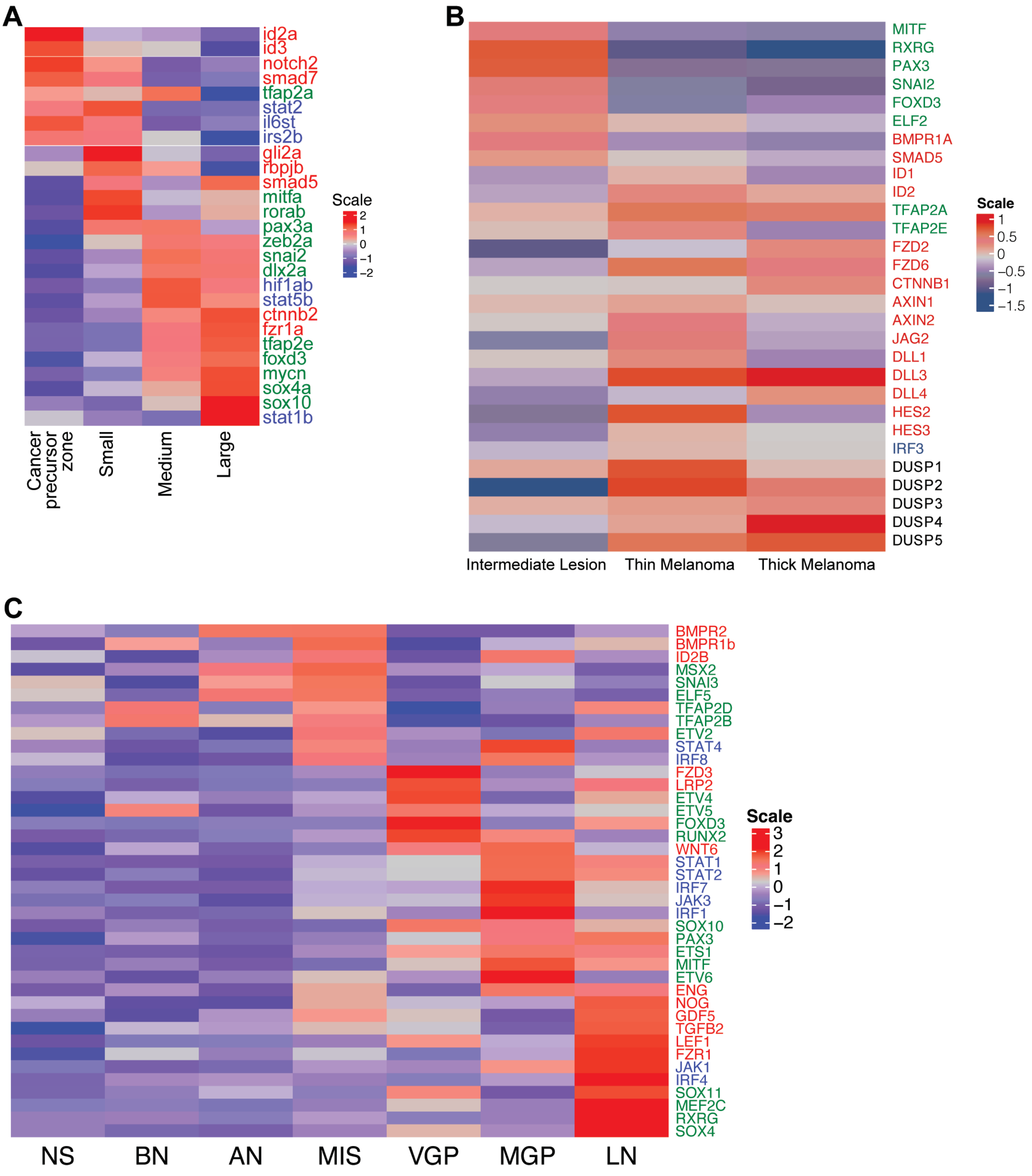

**Supplemental Figure 8 | Stage-specific gene expression during melanoma initiation in zebrafish and humans. A**, Gene expression of stage-specific differentially expressed genes from FACS-sorted melanocytes during melanoma initiation. **B**, Stage-specific expression signatures in intermediate lesions, thin melanoma, and thick melanoma samples taken from human patients from Shain et al. Cancer Cell, 2018. **C**, Stage-specific gene expression in human melanoma samples from microarray analysis performed in Smith et al. Cancer Biol Ther 2005. Samples were taken from normal skin (NS), benign nevi (BN), atypical nevi (AN), melanoma in situ (MIS), vertical growth phase (VGP), metastatic growth phase (MGP), and lymph node metastasis (LN).

### Supplemental Figure 9

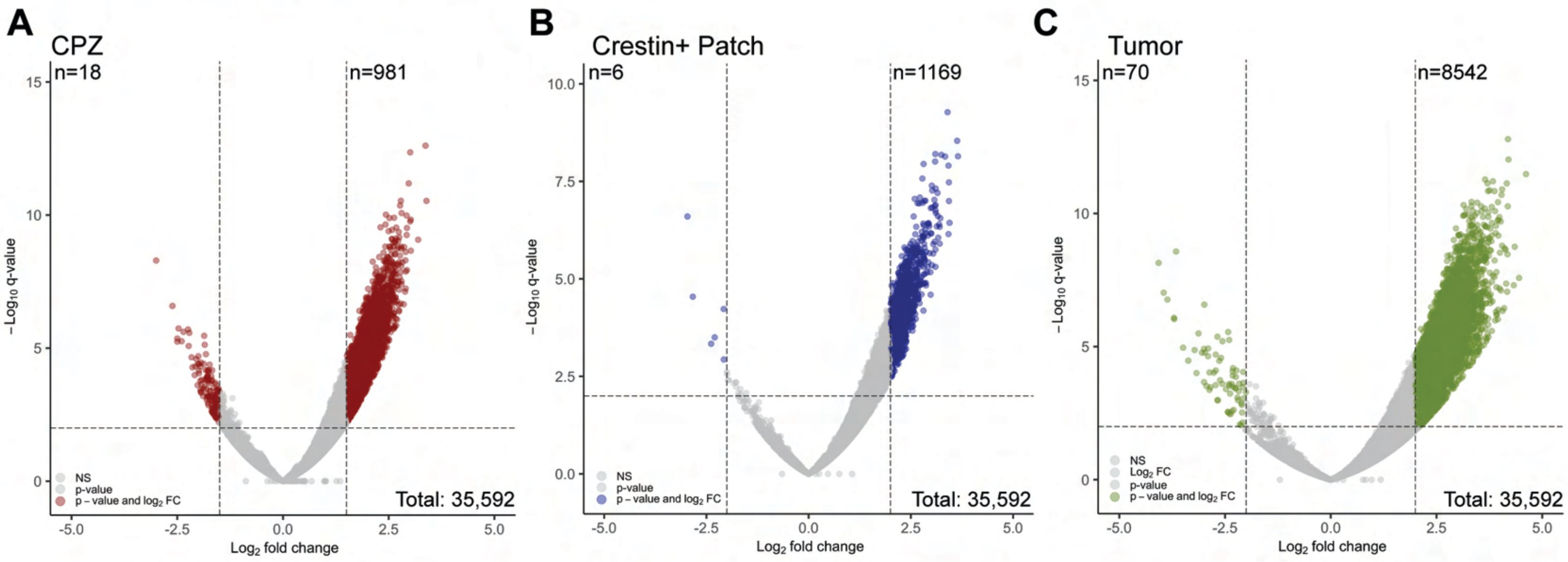

**Supplemental Figure 9 | Differential accessible chromatin regions across melanoma stages. A-C**, Volcano plots showing differentially accessible regions in CPZ (**A**), patch (**B**), and tumor (**C**) relative to cancerized field control. Dashed lines represent a 2 $log_2$FC (over cancerized field) and a 10e-3 q-value as assigned by DeSeq2. Colored regions indicate peaks that meet the logFC and q-value threshold.

### Supplemental Figure 10

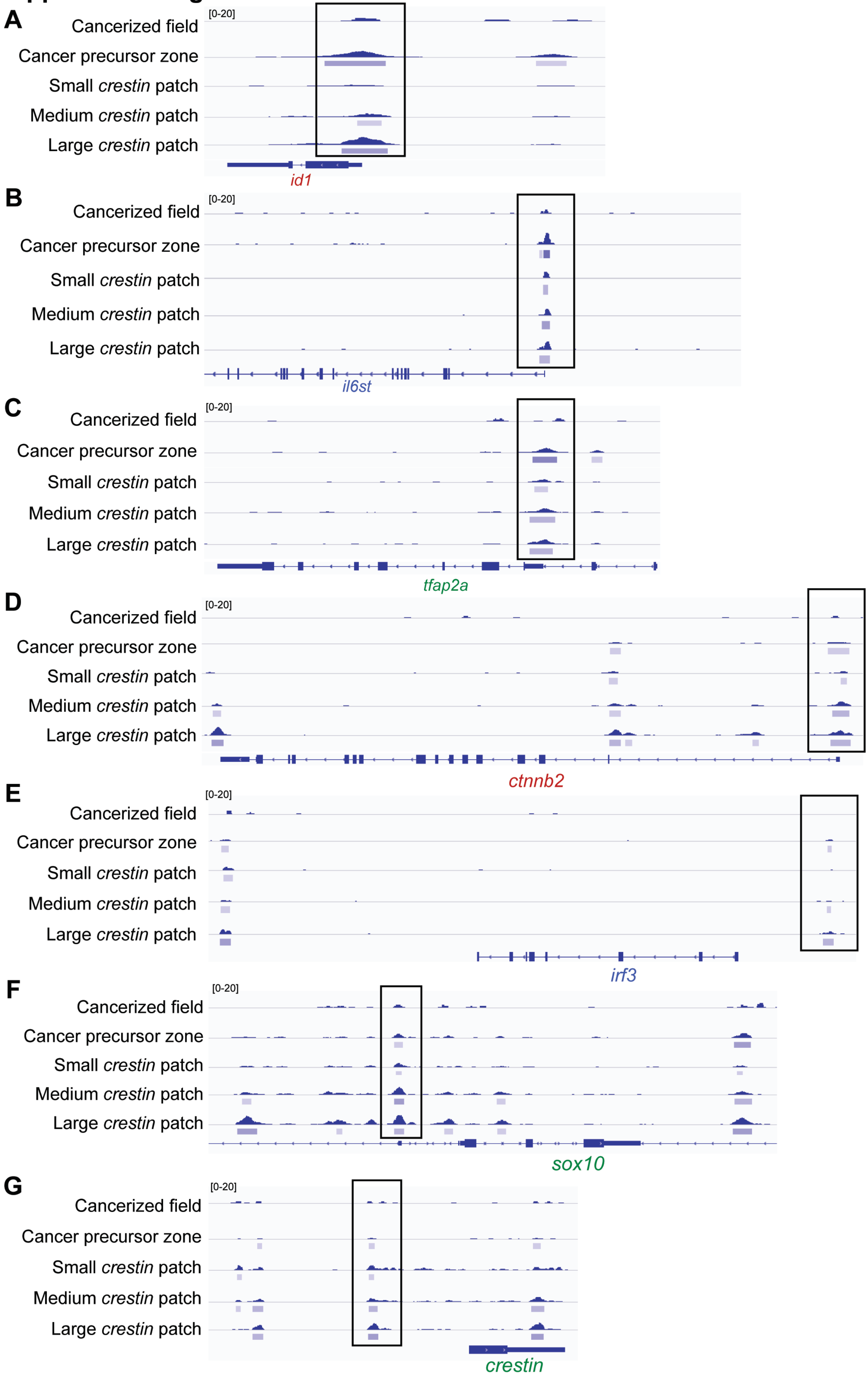

**Supplemental Figure 10 | Enriched binding motifs during melanoma initiation. A-G**, ATAC-seq peak tracks showing open chromatin in enhancers and/or promoters for the cancer precursor zone genes *id1* (**A**), *il6st* (**B**), and *tfap2a* (**C**), as well as the *crestin* patch genes *ctnnb2* (**D**), *irf3* (**E**), *sox10* (**F**), and *crestin* (**G**). Black box highlights the peak that is displayed in Fig. 2K.

### Supplemental Figure 11

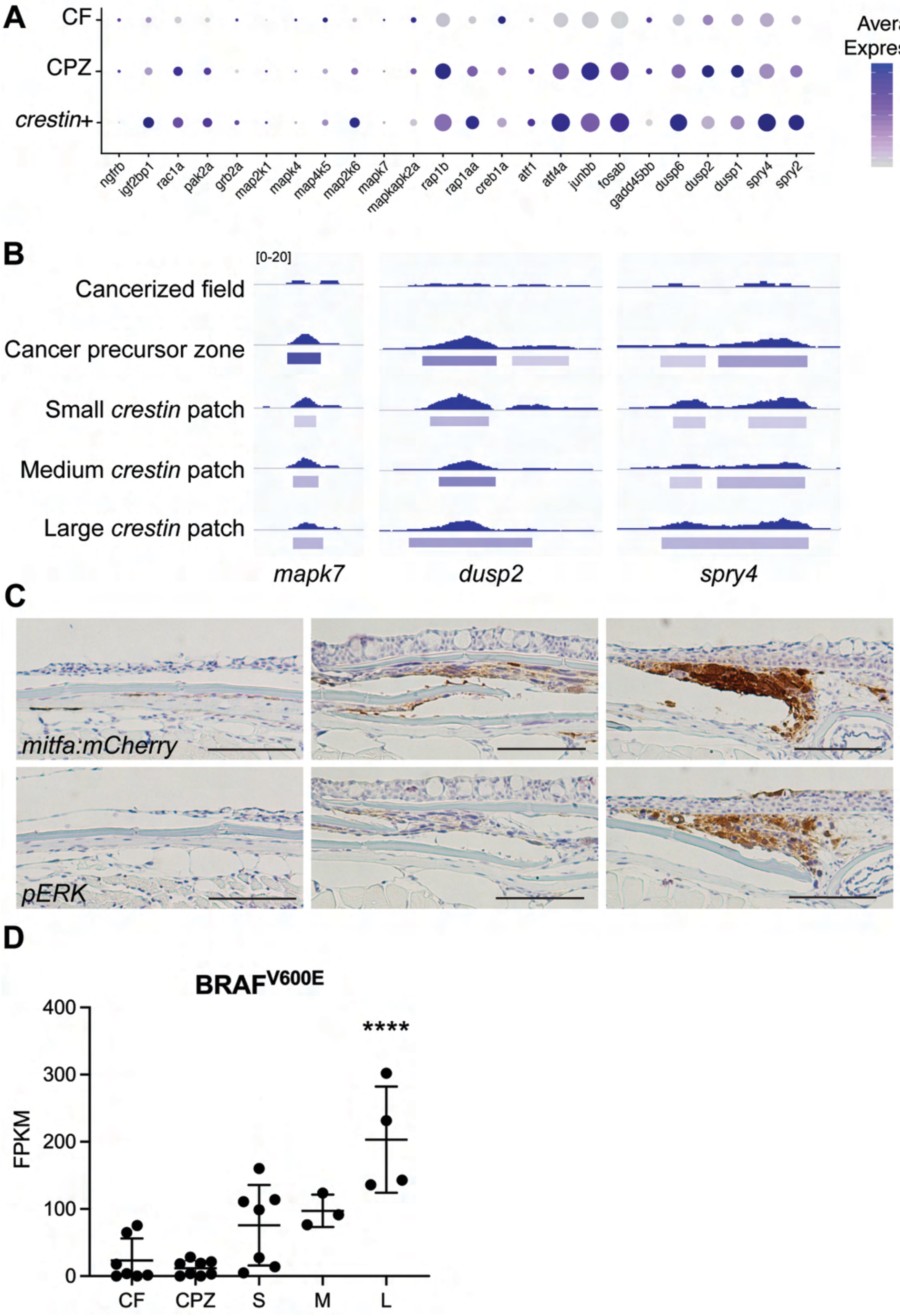

**Supplemental Figure 11 | MAPK pathway is not transcriptionally active in cancerized field melanocytes. A**, MAPK pathway gene expression. CPZ = cancer precursor zone. **B**, ATAC-seq peak tracks demonstrating that chromatin around MAPK pathway genes does not open until the formation of a cancer precursor zone. **C**, mCherry and phospho-ERK in cancerized field (left), early *mitfa* $^{med}$ cancer precursor zone (middle), and *mitfa* $^{high}$ cancer precursor zone (right). Scale bars, 100 μm. **D**, Expression levels of $BRAF^{V600E}$ are unchanged between FACS-isolated cancerized field (CF) and cancer precursor zone (CPZ) melanocytes, n=3-8. **** p≤0.0001.

#### Supplemental Figure 12

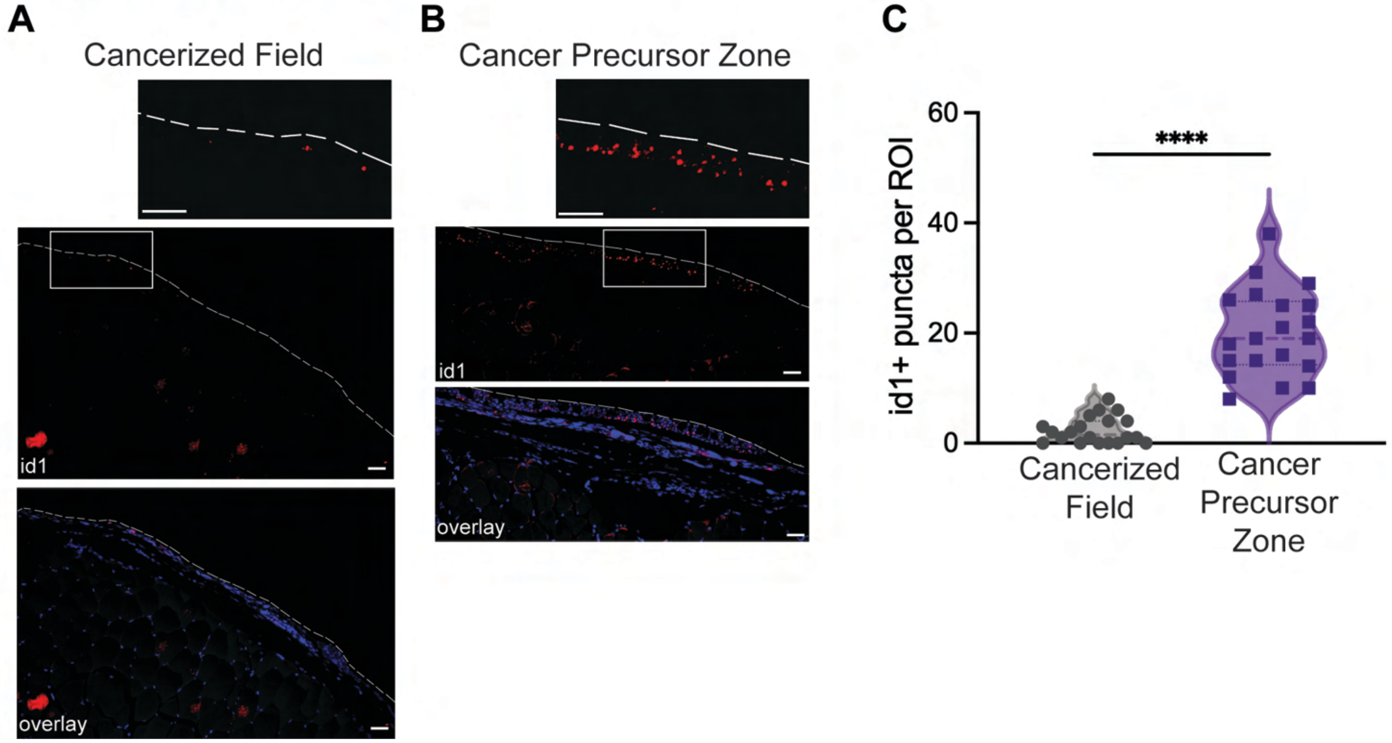

**Supplemental Figure 12 | ID1 expressed in zebrafish cancer precursor zones. A,B.** RNAScope in situ hybridization for id1 (red) and DAPI nuclear stain in fixed and sectioned cancerized field (**A**) or CPZ (**B**) melanocytes. Dashed line outlines the skin surface. **C**, Violin plot quantifying id1+ puncta per ROI in cancerized field and CPZ sections (n=4 sections; n=5 quantified ROI per section). ****, $p \leq 0.0001$.

### Supplemental Figure 13

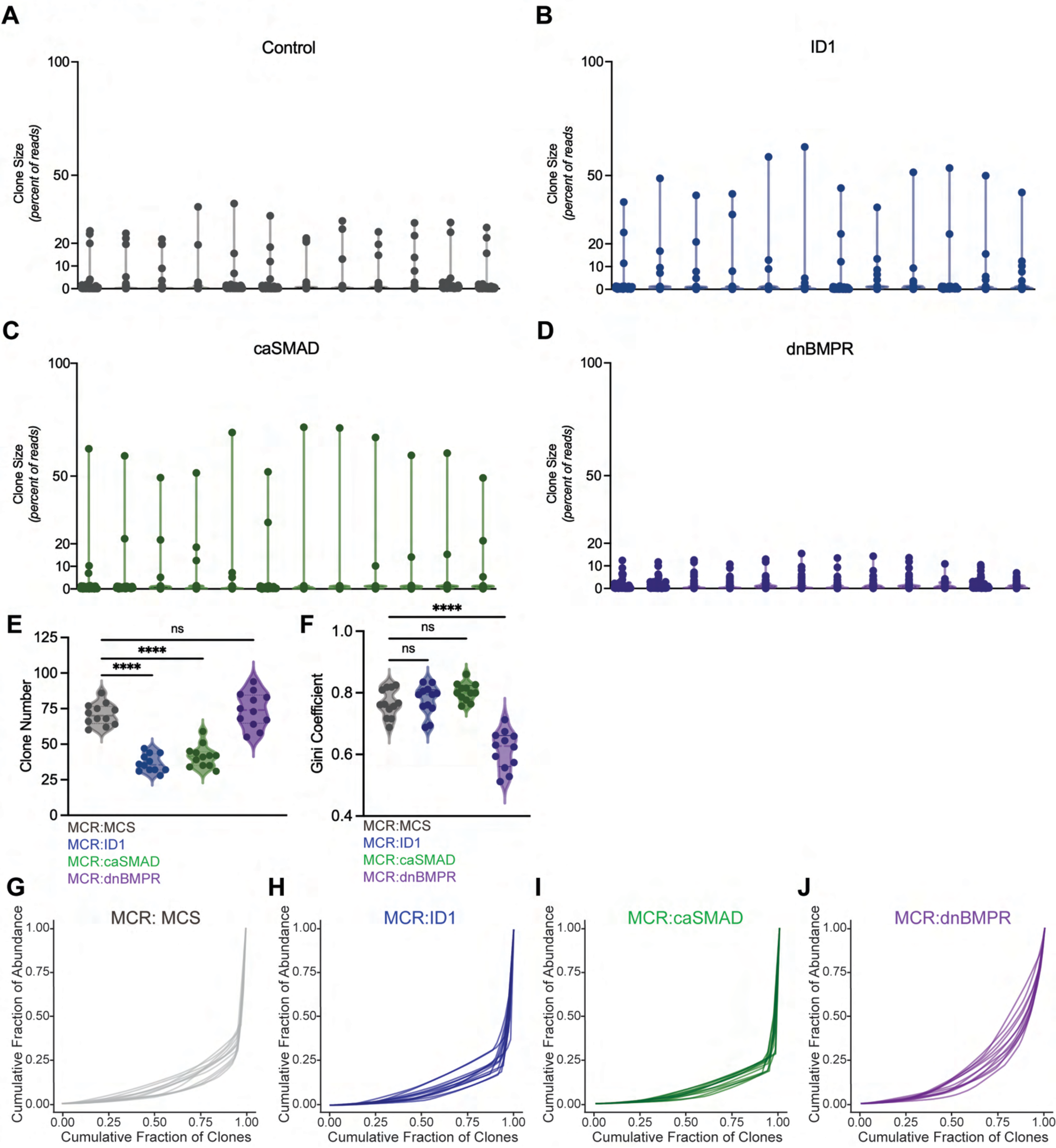

**Supplemental Figure 13 | BMP signaling modulates clonal dynamics of cancer precursor zones. A-D,** Violin plot of clone size (calculated by percent of reads) from individual GESTALT barcoded fish with MCR:MCS control (**A**), MCR:ID1 (**B**), MCR:caSMAD (**C**), and MCR:dnBMPR (**D**) melanocytes from Fig. 2E-G. Each dot is a clone and each line is a fish. **E**, Violin plot of total clone number (**E**) and Gini coefficient (**F**) (n=12 fish). n.s., not significant; ****, p≤0.0001. **G-J**, Lorenz curves of cumulative fraction of abundance and fraction of clones for control (**G**), ID1 (**H**), caSMAD (**I**), and dnBMPR (**J**) overexpressing CPZ melanocytes.

#### Supplemental Figure 14

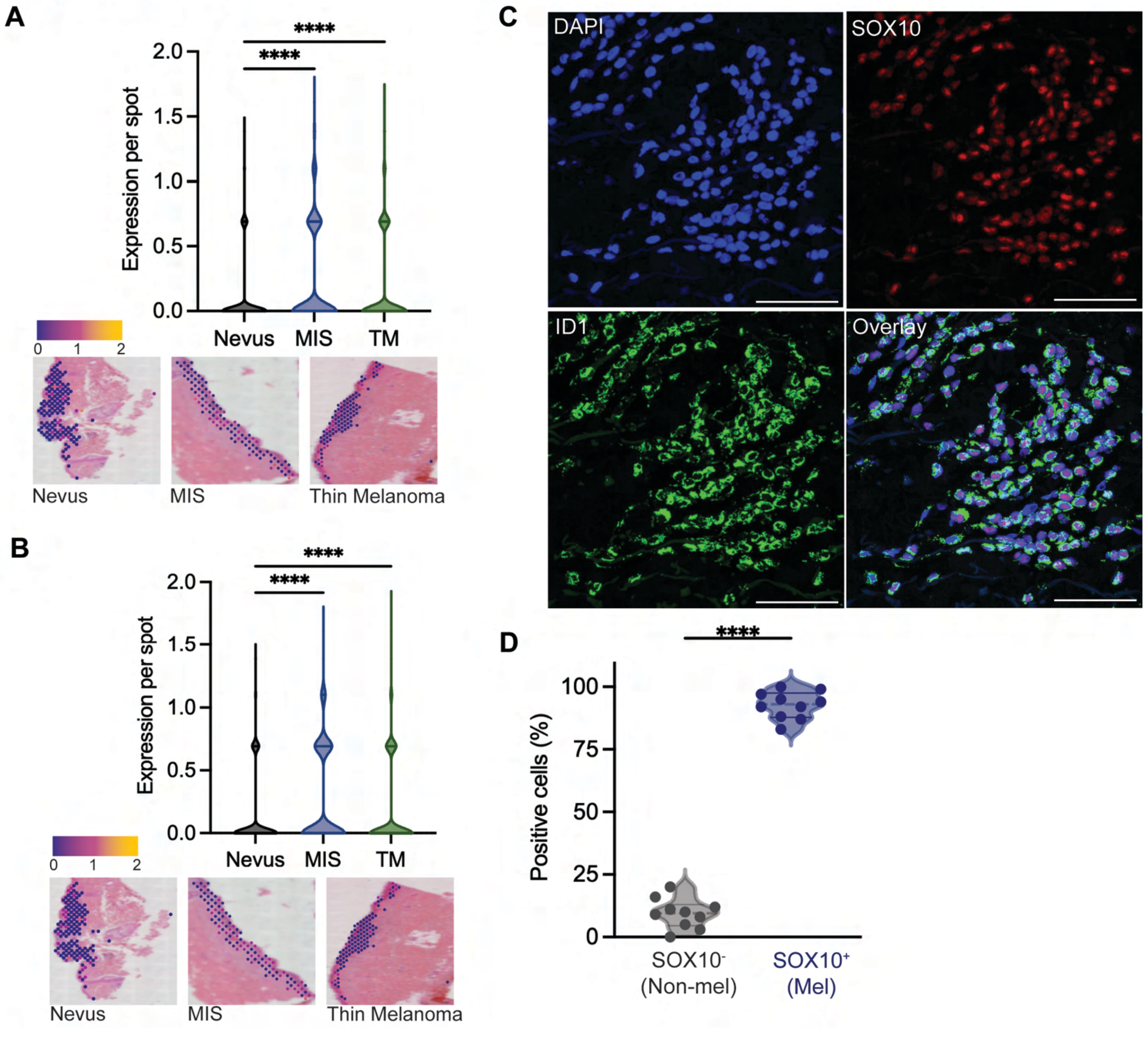

**Supplemental Figure 14 | ID1 is expressed in human melanoma in a stage-specific manner**. **A,B,** Spatial transcriptomic analysis of ID1 (**A**) and ID2 (**B**) in human nevi (n=1382 spots), melanoma in situ (MIS, n=642 spots), and thin melanoma (TM, n=1684 spots). ****, p≤0.0001. **C,** Immunofluorescence image of human nevi stained with DAPI (nuclei, blue), SOX10 (melanocyte, red), ID1 (green), and overlay. Scale bars, 50μm. **D,** Quantification of ID1 staining in SOX10+ melanocytes and SOX10- non-melanocyte across 10 human nevi sections. ****, p≤0.0001.

#### Supplemental Figure 15

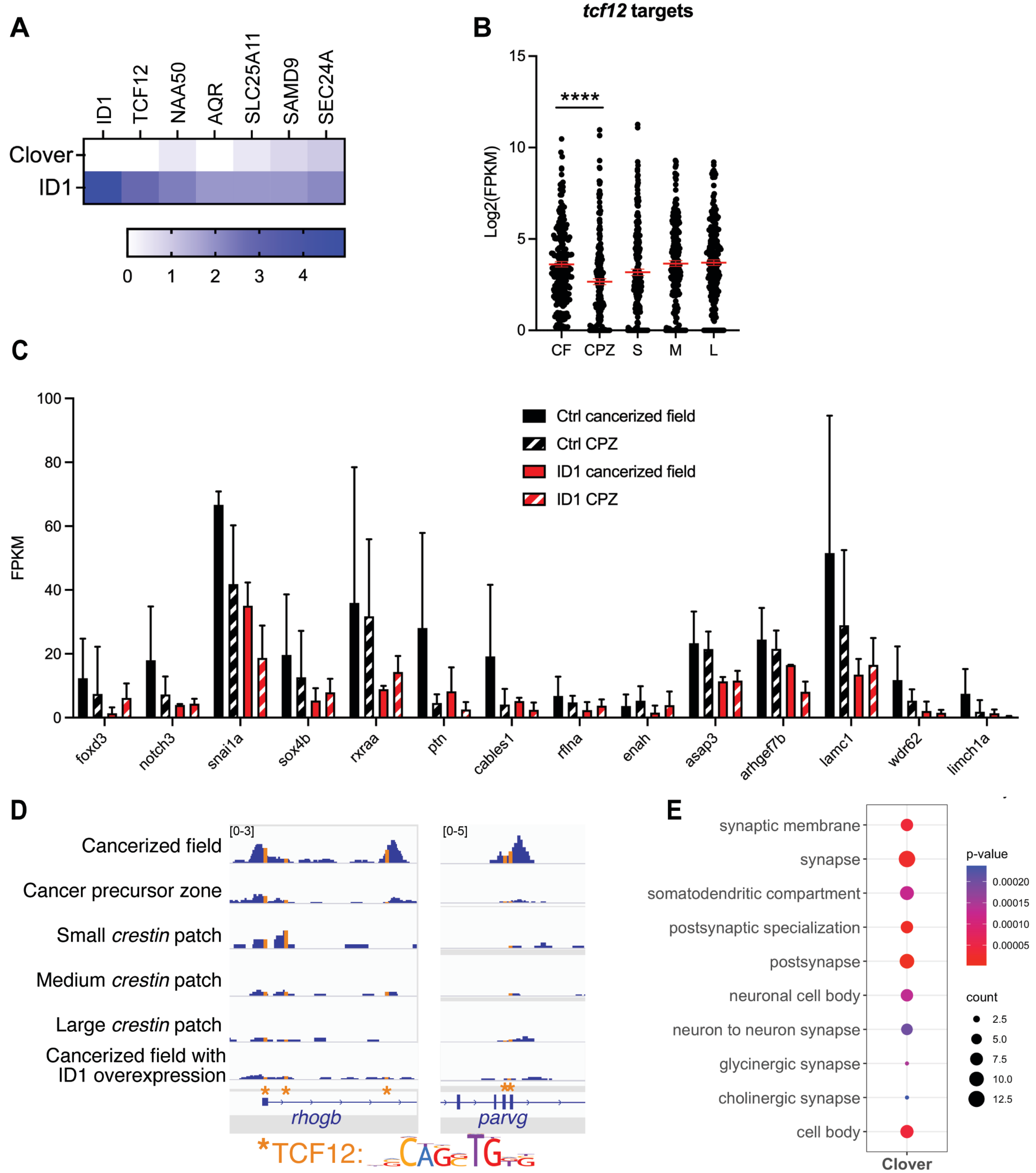

**Supplemental Figure 15 | ID1 inhibits TCF12, preventing binding to downstream targets and enabling cancer precursor zone formation. A**, Heatmap of the Log2FC of ID1-bound proteins over control. **B,** Expression of known TCF12 targets identified by the TRANSFAC Curated Transcription Factor Targets dataset. **C**, Expression of TCF12 targets decrease in cancer precursor zone and further decrease when ID1 is overexpressed. **D,** Increasing ID1 expression decreases open chromatin peaks in promoter or enhancer regions of *rhogb* and *parvg*. **E,** Dot plot of enriched GO terms in genes with annotated TCF12 motif lost following ID1 overexpression.
